# An ancient origin of the naked grains of maize

**DOI:** 10.1101/2024.12.02.626434

**Authors:** Regina Fairbanks, Jeffrey Ross-Ibarra

## Abstract

Adaptation to novel environments requires genetic variation, which may either predate the novel environment or arise as new mutations. The relative importance of standing genetic variation vs. *de novo* mutations in adaptation remains a fundamental question in evolutionary biology. Selection during domestication has been long used as a model to understand evolutionary processes, providing information not only on the phenotypes selected but also, in many cases, an understanding of the causal loci. Of the multiple causal loci that have been identified in maize, the selected allele can be found segregating in natural populations, consistent with their origin as standing genetic variation. The sole exception to this pattern is the well-characterized domestication locus *tga1*, which has long been thought to be an example of selection on a *de novo* mutation. Here, we use a large dataset of maize and teosinte genomes to reconstruct the origin and evolutionary history of *tga1*. We first estimated the age of *tga1-maize* using a genealogy-based method, finding that the allele arose approximately 41,000-49,000 years ago, predating the beginning of maize domestication. We also identify, for the first time, *tga1-maize* in teosinte populations, indicating the allele can survive in the wild. Finally, we compare observed patterns of haplotype structure and mutational age distributions near *tga1* with simulations, finding that patterns near *tga1* in maize better resemble those generated under simulated selective sweeps on standing variation. These multiple lines of evidence suggest that maize domestication likely drew upon standing genetic variation at *tga1* and cement the importance of standing variation in driving adaptation during domestication.

## Introduction

Adaptation to novel environments requires genetic variation, which may either predate the novel environment or arise as new mutations. Increasingly, case studies reveal significant contributions of pre-existing (standing) genetic variation to adaptation in both experimental (Teotónio *et al*. 2009; Burke *et al*. 2014; Sheng et al. 2015) and natural populations (Pennings 2012; Lai et al. 2019; Chaturvedi et al. 2021). However, the relative importance of standing genetic variation vs. *de novo* mutations in adaptation, and the factors shaping that balance, remain fundamental questions in evolutionary biology. To address this question, the genetic variants underlying adaptive phenotypes must first be identified, then categorized as either standing variants or *de novo* mutations. Typically, the latter step involves comparisons between closely related populations or with past samples from the same population. Though challenging in most systems, the extensive genomic resources and well-defined phenotypes of domesticated crops provide a tractable model for studying the genomic basis of adaptation (Ross-Ibarra *et al*. 2007).

Maize domestication has proven a particularly fruitful area of exploration for evolutionary biologists. Comparisons between maize and its extant wild relatives, as well as between modern and archaeological maize, have yielded insight into the phenotypes under selection during maize domestication (Doebley 2004). Though the identification of causal variants themselves remains elusive, a handful of well-characterized loci provide an opportunity to assess the relative contribution of standing genetic variation and *de novo* mutations in adaptation during domestication. For most of these well-characterized loci, including *tb1* (Studer et al. 2011), *gt1* (Whipple et al. 2011), *KRN4* (Liu et al. 2015), and *ZmSh1-1* (Lin et al. 2012), evidence suggests that maize domestication primarily leveraged standing genetic variation. The causal allele at these loci can be found segregating in modern wild populations, suggesting these alleles arose before domesticated and wild populations split.

The preeminence of standing genetic variation in maize domestication is ostensibly contradicted by the well-known domestication locus, *tga1* (*teosinte glume architecture1*). *Tga1* largely controls the most critical step in maize domestication: the loss of the kernel fruitcase (Dorweiler et al. 1993; Wang et al. 2005). In the wild relative, teosinte, a fruitcase tightly encloses each individual kernel (Figure 1A). Thought to protect the plant from seed predators, the fruitcase also makes teosinte kernels extraordinarily difficult to process for human consumption. During domestication, selection upon a single nonsynonymous G to C mutation in *tga1* contributed to the evolution of the fully exposed kernels seen in maize today (Wang et al. 2015, hereafter we refer to the G and C alleles as *tga1-teosinte* and *tga1-maize*, respectively). Unlike other well-characterized domestication loci, however, both field surveys (Iltis 2006) and population genetic studies (Wang et al. 2005, 2015) report an absence of both the phenotype and its causal variant in the wild. Many argue that *tga1-maize*, should it appear in the wild, would be too deleterious to survive beyond just a few years due to the likely increased susceptibility of exposed kernels to pests, predators, and pathogens. Under cultivation, however, the benefits of exposed kernels apparently outweigh the risks, as estimates suggest strong selection for exposed kernels during domestication (Wang et al. 2005). Given these findings, the researchers who first identified and described *tga1-maize* argue that exposed kernels, and the underlying genetic variant *tga1-maize*, arose *de novo* during domestication.

**Figure 1.**
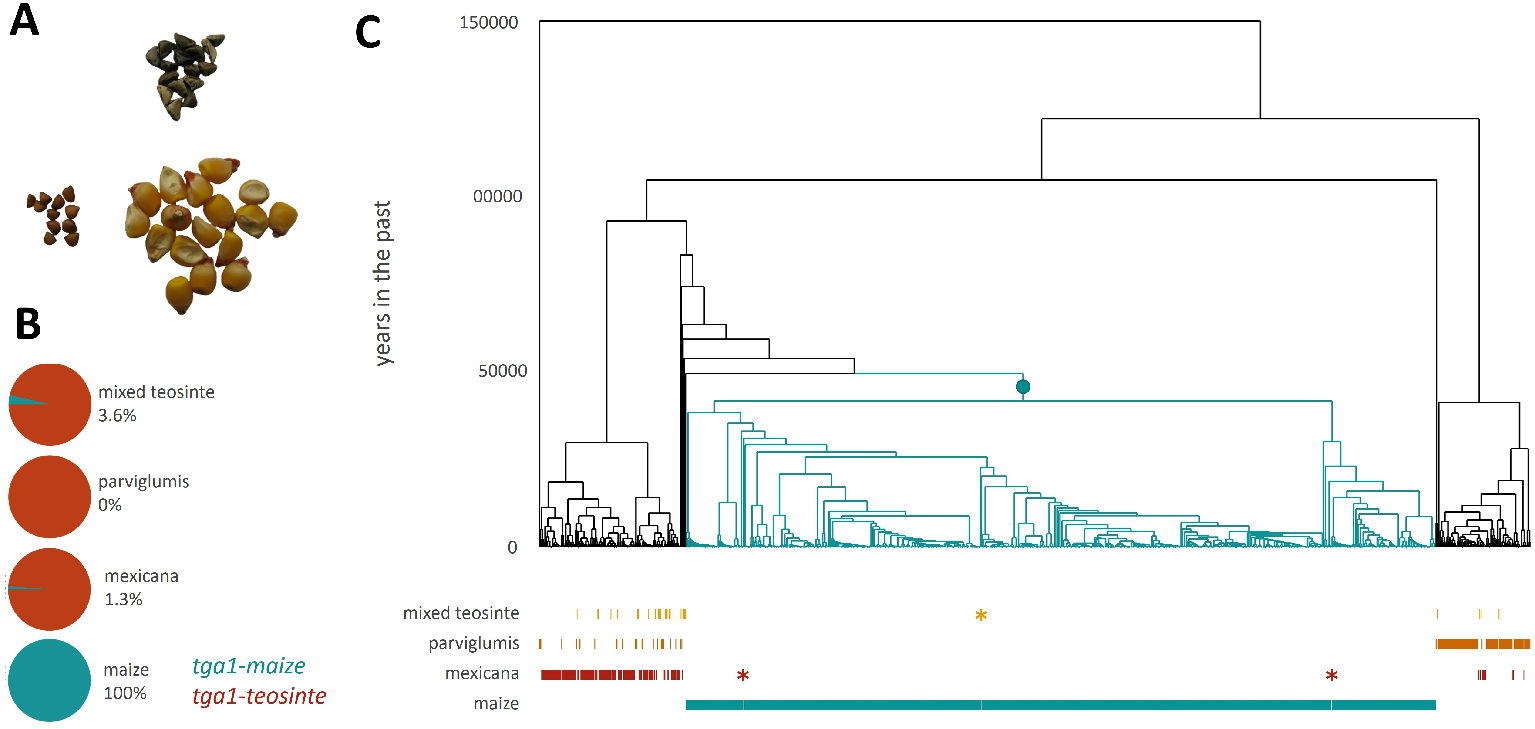
*Tga1-maize* predates the onset of domestication. *(A)* Teosinte kernels with (top right) and without fruitcases (bottom left); maize kernels (bottom right). *(B)* Shown is the marginal genealogy at *tga1*, with the *tga1-maize* allele mapped onto the tree as a blue circle on a branch spanning 41,586 to 49,476 years in the past. The population of origin of each tip is shown below the genealogy, with copies of *tga1-maize* found in teosinte populations highlighted with asterisks. *(C)* Pie charts show the frequency of the *tga1-maize* allele in each group.

Yet, recent population genetic work casts doubt on this simple model for exposed kernels and *tga1-maize*. Under classic population genetic theory, strong, recent selection on *de novo* mutations should produce characteristic signatures in the genome (i.e., reduced diversity, elevated linkage disequilibrium, and shift toward excess rare and high frequency derived variants). Despite this, and *tga1*’s inarguable importance for domestication, genome-wide scans for signatures of selection do not consistently identify *tga1* as one of the most prominent candidate genes (Hufford *et al*. 2012; Xu *et al*. 2022).

Driven by these conflicting avenues of evidence, in this paper we reevaluate the domestication history of *tga1*. Given the extremely limited archaeological record before and during maize domestication, current archaeological data cannot yield direct information about when exposed kernels or associated alleles arose. Instead, we turned to population genetics to reconstruct the origin and evolutionary history of *tga1*. Leveraging a large dataset of maize and teosinte genomes, we used a genealogy-based approach to estimate the age of *tga1-maize*. We estimated that the causal mutation arose approximately 41,000-49,000 years ago, predating even conservatively early dates for the beginning of maize domestication. We then used this same dataset to identify, for the first time, *tga1-maize* in wild teosinte populations. Finally, we compare observed patterns of haplotype structure and mutational age distributions near *tga1* with simulated selective sweeps. We find that observed patterns near *tga1* in maize better resemble those generated under simulated selective sweeps on standing variation, rather than on *de novo* mutations. These multiple lines of evidence suggest that, in contrast to decades of thought based on the absence of evidence, domestication likely drew upon standing genetic variation at *tga1*. This work contributes to growing evidence for a fundamental role of standing genetic variation in both maize domestication and adaptation more broadly.

## Results

### The causal tga1 mutation exists in modern teosinte

If *tga1-maize* first arose in wild teosinte populations before domestication began, we hypothesized it should be observable in a sufficiently large sample of wild teosinte. Previous work analyzing relatively small numbers of teosinte genomes (n = 12 sampled alleles in Wang *et al*. (2005) and n = 20 alleles in Wang *et al*. (2015)) failed to identify *tga1-maize* in wild *parviglumis* populations, reporting instead that *tga1-maize* is the single fixed difference between maize and teosinte in *tga1*. We questioned whether the fixed difference would hold with substantially larger samples of teosinte genomes from maize’s closest relatives, *Zea mays subsp. parviglumis* and *Zea mays subsp. mexicana* (n = 328 sampled alleles, including n = 140 *parviglumis* alleles). In agreement with previous work, we did not find *tga1-maize* in modern *parviglumis* genomes. We did, however, identify the allele in two *mexicana* individuals (samples 5A2 and 6A12) and one mixed teosinte individual (5G4). All three individuals were heterozygous, carrying one *tga1-teosinte* allele and one *tga1-maize* allele. The *tga1-maize* allele therefore appears at a frequency of 1% (3 out of 328 haplotypes) in combined teosinte samples or 1.25% (2 out of 160 haplotypes) when considering the subspecies *mexicana* alone (Figure 1B). *Tga1-maize* has not previously been reported in any wild teosinte genome.

### tga1-maize predates domestication

Though it remains unclear when the *tga1-maize* mutation first arose, population genetic evidence indicates it may have been fixed in maize ≈ 10, 000 years ago (Wang *et al*. 2005). Archaeological data support the idea that *tga1-maize* may have fixed early during maize domestication, as the earliest archaeological maize cobs exhibit the exposed kernel phenotype (Piperno and Flannery 2001) and the *tga1-maize* allele was found in the oldest ancient genome sampled to date (Ramos-Madrigal *et al*. 2016).

Given its observation in modern teosinte samples, we sought to estimate the age of *tga1-maize*. We used Relate (Speidel et al. 2019) to estimate the ancestral recombination graph for a 5Mb region centered on *tga1* from whole genome sequencing data for 507 modern maize (*Zea mays subsp. mays*) lines and 164 teosinte individuals (Figure 1). Our teosinte samples include the two subspecies most closely related to maize, namely 70 *parviglumis* individuals, 80 *mexicana* individuals, and 14 teosinte individuals of mixed ancestry (*Zea mays* mixed). While domesticated maize originally evolved from ancestral *parviglumis* populations, later introgression with *mexicana* contributed to maize diversification and dispersal (Yang et al. 2023). We mapped the *tga1maize* mutation onto the marginal genealogy reconstructed at the causal nucleotide, allowing us to estimate the age of the causal mutation as the midpoint of the branch upon which it arose. We obtained an estimate of ≈ 45, 531 years in the past, placing the *tga1-maize* mutation far earlier than accepted estimates for the onset of domestication of ≈ 9, 000 − 15, 000 years ago (Beissinger et al. 2016; Piperno et al. 2009) and earlier even than the likely first human occupation of the Americas (Dillehay et al. 2015).

### The tga1-maize persists in modern teosinte populations

If the observed *tga1-maize* copies found in modern teosinte are examples of segregating ancestral standing variation, we might expect relatively deep coalescence between the maize and teosinte copies of the allele. Instead, our estimated ARG reveals very short branches subtending maize-teosinte coalescences, suggesting a relatively recent origin of the *tga1-maize* copies in modern teosinte.

Extensive evidence for widespread historical and ongoing gene flow between maize and teosinte (Ross-Ibarra et al. 2009; Fukunaga *et al*. 2005; Moreno-Letelier et al. 2020) raise the possibility of an introgressed origin for the *tga1maize* allele observed in modern teosinte. We estimated maize and teosinte ancestry along the genomes of each of the heterozygous samples using ELAI (Guan 2014, ; Figure 2). Estimates of maize ancestry for two of the samples, the *mexicana* 5A2 and the mixed teosinte 5G4, are largely consistent with previous estimates of nearby wild populations (2.7% for 5A2 compared to an average of 4.6% in the San Pedro population 7 km distant and 32.0% of 5G4 compared to an average of 32.8% in the nearby Nabogame population 36km away; Calfee *et al*. (2021)). These similarities to population-wide averages suggest the samples are not recent hybrids, but instead reflective of overall levels of admixture in the wild populations from which they were sampled. In contrast, the *mexicana* sample 6A12 carries a greater proportion of maize ancestry, estimated at 15.4%, than the nearby Malinalco population 26 km away, where sampled *mexicana* individuals possess a maximum of 2.2% maize ancestry.

**Figure 2.**
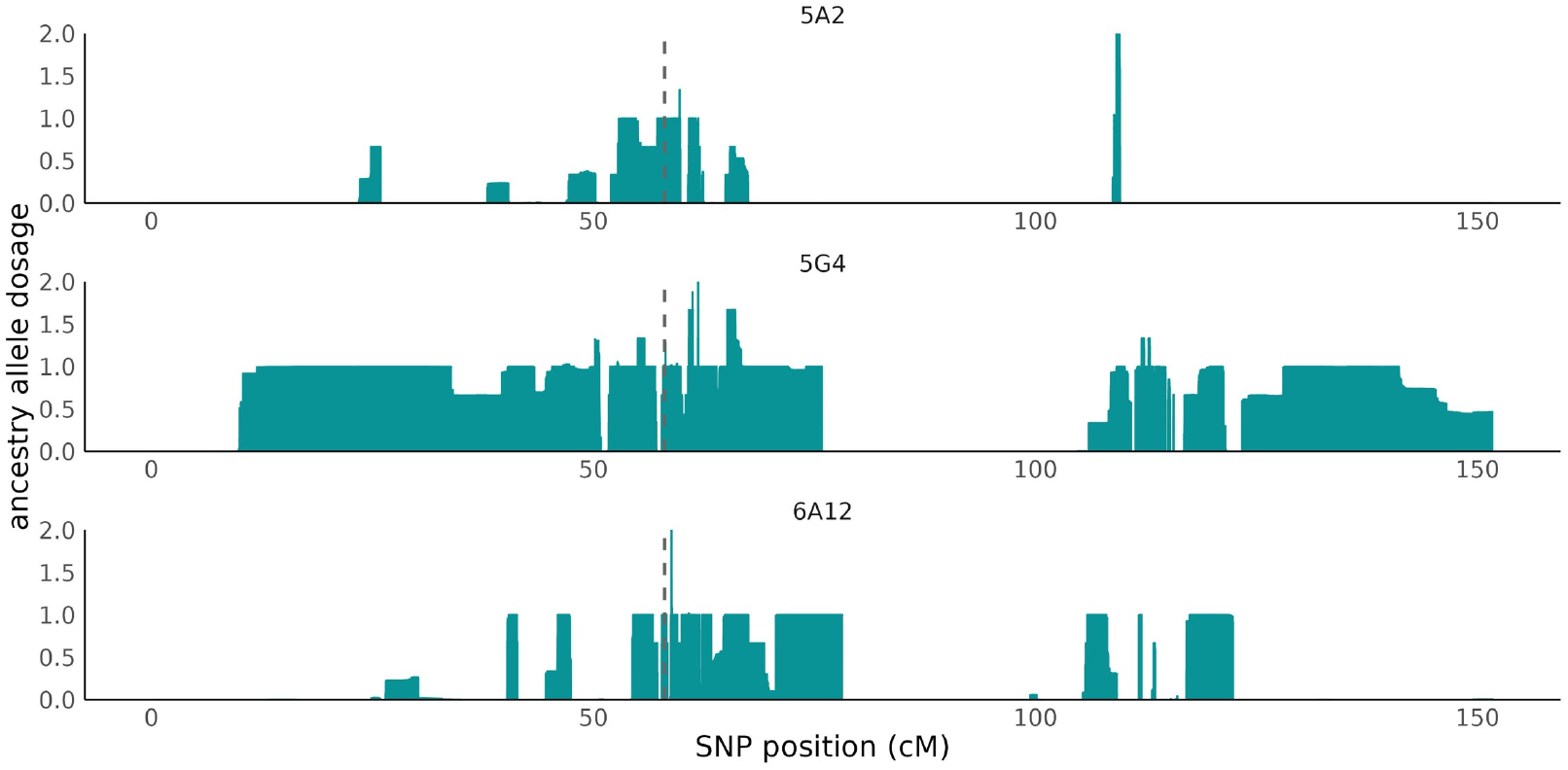
*tga1-maize* in teosinte is the result of gene flow from modern maize. ELAI estimates for maize ancestry across chromosome 4 in the three heterozygous teosinte individuals. Shown at each position is the mean estimated number of maize alleles across three independent runs per individual. The dashed line indicates the causal site position.

If *tga1-maize* recently entered teosinte populations via gene flow, then we would expect to observe an extended maize-like haplotype spanning the *tga1* locus. A relatively old introduction, however, should generate a short, localized maize-like haplotype and may not be distinguishable from patterns expected if *tga1-maize* arose within teosinte itself. Ongoing gene flow between *mexicana* and maize (Calfee *et al*. 2021) complicates efforts to date hybridization, but a simple estimate based on the genetic length of the introgressed haplotypes suggests that the observed *tga1-maize* allele was introduced ≈ 100, ≈ 140, and ≈ 180 years ago for 5A2, 5G4, and 6A12, respectively. Although the inferred times are recent compared to the timing of maize domestication and suggest *tga1-maize* did not originate in any of these wild populations, these times nonetheless suggest that *tga1-maize* may persist in wild teosinte populations for many years and survive in multiple, diverse environments.

### Haplotype structure and nearby mutational age distributions at tga1 resemble those generated by selection on standing genetic variation

Previous work reported at least two common haplotypes at *tga1* in maize (Brandenburg *et al*. 2017), which we confirmed by clustering and visualizing genetic diversity in a sample of 507 inbred maize lines across a 10 Kb window encompassing *tga1* (Figure S4). These findings suggest that the haplotype structure at *tga1* differs from that expected if *tga1-maize* arose as a *de novo* mutation, which should generate a single predominant, fixed haplotype under classical population genetic models (Garud et al. 2015). To more explicitly evaluate whether a standing genetic variation origin for *tga1-maize* can explain nearby haplotype structure in modern maize, we conducted population genetic simulations in SLiM version 4.0.1 (Haller and Messer 2023) and msprime (Kelleher et al. 2018). We first used a simple population bottleneck model previously estimated for maize (Beissinger *et al*. 2016). Under this demographic model, maize experienced an instantaneous bottleneck approximately ≈ 15,500 years ago, followed by exponential population growth until the present. For *de novo* simulations, we introduced a new mutation immediately after the population bottleneck, corresponding to the beginning of domestication. For standing genetic variation simulations, we randomly selected a neutral mutation of a given frequency and switched its selection coefficient to 0.05, consistent with previous estimates for *tga1* of 3-4% (Wang *et al*. 2005). We also specified a locally elevated recombination rate estimated with LDHelmet (Figure S5).

We first found that, as expected, selected standing variants fixed at higher proportions than *de novo* mutations for all tested selection coefficients. Indeed, for this simple bottleneck demographic model, *de novo* mutations only fix in about one-third of simulations even at the strongest selection coefficient (Table S2).

To compare the haplotype diversity patterns, we used 50-SNP sliding windows to calculate the haplotype homozygosity statistics H12 and H2/H1 (Garud *et al*. 2015) on both the empirical data at *tga1* from 507 maize inbred genomes and the simulated selective sweep data. H12 estimates the combined haplotype homozygosity for the first and second most frequent haplotypes in a window, thereby capturing overall signals of selection. H2/H1 evaluates the ratio of H2, the haplotype homozygosity calculated using all but the most frequent haplotype within the population, to H1, the haplotype homozygosity of the most frequent haplotype. High H2/H1 values indicate elevated haplotype diversity, expected to be generated by selection on standing variation, while low H2/H1 values represent low haplotype diversity, expected to result from selection on *de novo* mutations.

For this simple bottleneck demographic model, *de novo* simulations consistently generate H2/H1 values lower than those observed in modern maize (Figure 3, top row; Figures S6 and S7, first column). In fact, only a model in which the causal mutation started at a frequency of 5% in the population at the beginning of domestication produces H2/H1 values substantially similar to the observed (Figure 3, top row). This result holds even when selection is weak (Figure S6), but is less robust to the details of the local genetic map (Figure S7). While these results are consistent with selection acting on standing genetic variation at *tga1*, we note that our simulations do not perfectly replicate patterns in the observed maize data, highlighting the limitations of relatively simple simulation scenarios.

**Figure 3.**
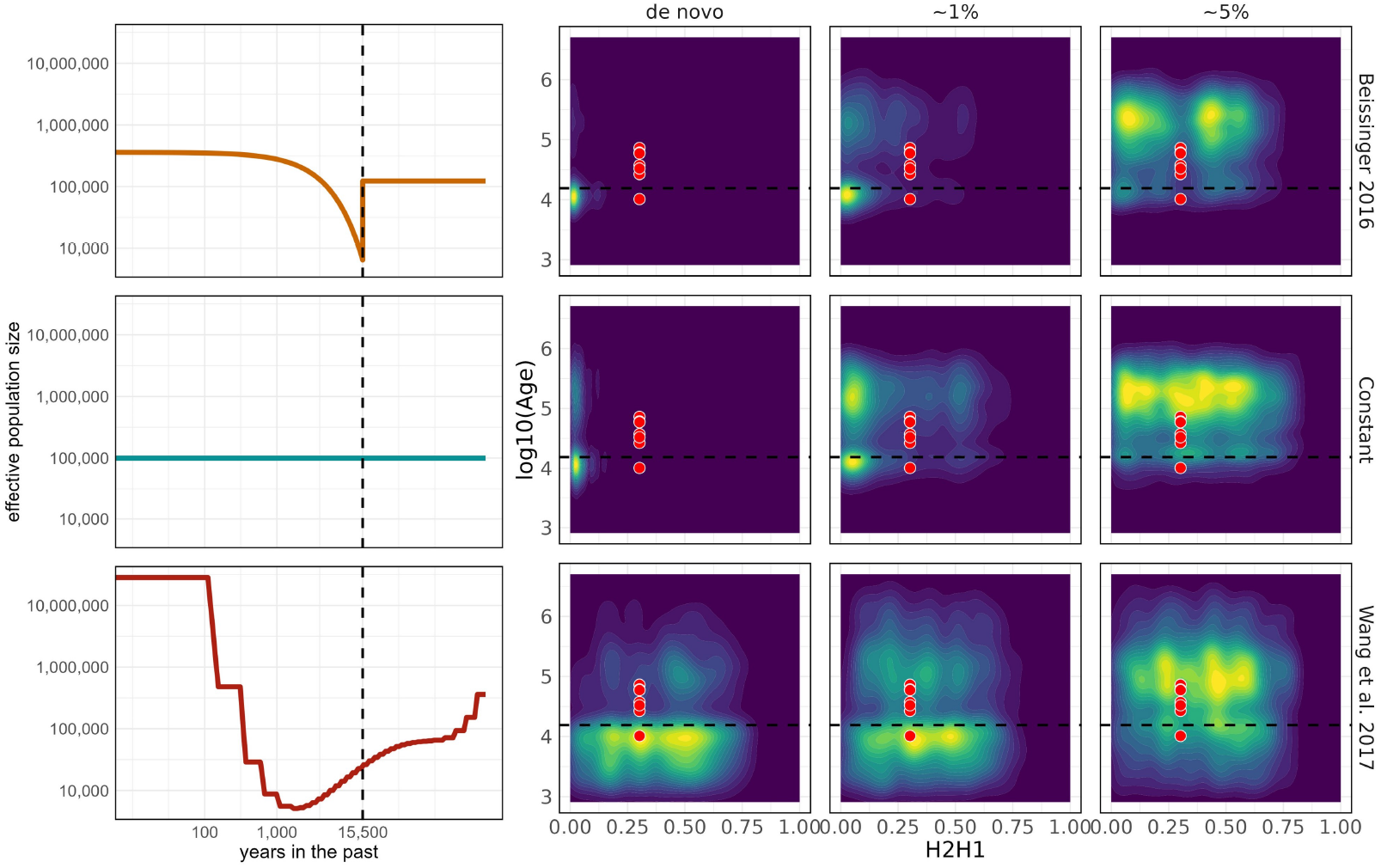
Haplotype structure and mutational age distributions near *tga1* resemble selection from standing genetic variation. Demographic models simulated in SLiM and msprime shown on left. Density plots represent the H2/H1 values and mutational ages from 300 simulation runs where mutation fixed for each combination of demographic model (rows) and starting frequency for the selected mutation (columns). In each plot colors range from dark blue (low density) to bright yellow (high density). Black dotted lines represent the onset of domestication/selection at 15,500 years in the past. Simulations were run with a selection coefficient of 0.05 and a recombination rate of 1e-7 (see Figure S8 for lower recombination rate). Red points indicate values observed in 50-SNP sliding windows centered around *tga1* in a sample of 507 modern maize inbreds.

The demography of maize domestication remains poorly understood, despite several attempts to infer reasonable demographic models for maize and teosinte (Beissinger et al. 2016; Wang et al. 2017; Wright et al. 2005; Eyre-Walker et al. 1998; Tenaillon *et al*. 2004). Because demography can confound signatures of selection upon standing genetic variation (Peter *et al*. 2012), we conducted additional simulations to assess whether our inferences were robust to different demographic models. In addition to the simple bottleneck used previously (adapted from Beissinger *et al*. (2016)), we performed simulations with the same range of parameters under both a constant size population and a complex demographic bottleneck inferred from pairwise coalescence rates (Wang et al. 2017).

Simulations performed under the model with constant population size largely mirror our previous results (Figure 3, second row; Table S3). In contrast, both *de novo* and standing genetic variation simulations implemented with the complex bottleneck model produce H2/H1 values similar to observed *tga1* values (Figure 3; bottom row). In addition, we observe that *de novo* mutations fix at higher rates under the more complex demographic model, but still not as high as standing genetic variants (Table S4). The discrepancy likely arises because the Wang *et al*. (2017) model specifies a recent, prolonged bottleneck beginning thousands of years before domestication begins. As a result, under our model of *de novo* mutation with an origin ≈ 15,500 years ago, the mutation sweeps to fixation before the completion of the population bottleneck in the more complex model. During the period between the mutation’s fixation and the bottleneck, neutral mutations arise on the genetic background of the fixed, selected mutation. As a consequence of the bottleneck, a random sampling of these neutral haplotypes survive and then rise in frequency in the population, producing an excess of intermediate-frequency alleles and elevated H2/H1 values that may resemble a sweep from standing genetic variation. Further, the complex bottleneck model infers a more substantial population growth rate after the bottleneck, allowing for the accumulation of overall greater genetic diversity. Thus, under some demographic histories, selection on *de novo* mutations cannot be clearly differentiated from selection on standing genetic variation when evaluating haplotype structure alone.

To further differentiate between *de novo* and standing genetic variation models, we investigated the age distribution of mutations near the causal *tga1* site. After a selective sweep on a standing variant, multiple haplotypes will rise to intermediate frequency, such that the only fixed mutation within the sweep should be the selected mutation itself. As a result, many of the nearby mutations should predate the sweep. In contrast, after a selective sweep on a *de novo* mutation, both the *de novo* mutation and tightly linked, nearby mutations will rise to fixation. Nearby mutations still segregating in the population should primarily be those that arose during or after the sweep, resulting in a nearby mutational age distribution younger than the sweep itself. We compared the mutational ages estimated from our ancestral recombination graph at *tga1* to those of the simulated sweeps, finding that, under all demographic models, the observed mutational age distributions best match those generated under selection on standing variation (Figure 3, right two columns).

Combined, the observed H2/H1 and mutational age distributions near *tga1* best resemble those generated under selection on standing genetic variation (Figure 3).

## Discussion

We present evidence that overturns the widely accepted model of *tga1* evolution during maize domestication. Rather than the *de novo* appearance of *tga1-maize* representing a “eureka moment” that catalyzed maize domestication (Iltis 2004), we instead argue that *tga1-maize* existed as standing genetic variation in wild, ancestral teosinte populations. First, we estimate that the causative mutation in *tga1, tga1-maize*, predates the earliest evidence of domestication by up to ≈ 35,000 years. Further, we find the first clear evidence of the *tga1-maize* allele in teosinte. Our inference of the timing of gene flow suggests that, contrary to prior assumptions, *tga1-maize* may persist in wild teosinte populations beyond just a few years. Finally, we find that simulated selective sweeps from standing genetic variation, but not from *de novo* variation, produce both haplotype diversity patterns and mutational age distributions most similar to those near *tga1* in modern maize.

### Tga1 in the wild

Our age estimate for the causative mutation in *tga1* suggests that the allele arose in an ancestral teosinte population approximately 25,000-35,000 years before the onset of domestication from ancestral *parviglumis* populations. Yet neither our sampling nor previous work have identified the *tga1-maize* allele in wild *parviglumis* populations (Wang et al. 2005, 2015). One possible explanation for this apparent discrepancy is that the allele persists in modern *parviglumis* populations, but sampling strategies may hinder its detection. While previous efforts sampled a limited number of *parviglumis* individuals that fail to capture most rare alleles in a population, even our sample of 140 *parviglumis* alleles failed to identify *tga1-maize*. Spatial variation in the presence of *tga1-maize* could explain these results if *tga1-maize* exists at low frequencies in just one or a few *parviglumis* populations, but is absent in others. Consistent with this possibility, examination of short read data from Tittes *et al*. (2021) identifies the *tga1-maize* allele in a *parviglumis* population, but sequencing depth from these samples precludes confident genotype assignment. Further, extant teosinte populations exhibit substantial variation in the frequency of other domestication alleles. For example, the frequency of the maize allele of the canonical domestication locus *tb1* ranges wildly from absent up to 44% in different teosinte populations (Vann *et al*. 2015). Declines in and extinctions of some *parviglumis* populations associated with anthropogenic activities (Sánchez González *et al*. 2018; Rivera-Rodríguez *et al*. 2023) may have also caused the stochastic loss of populations harboring rare *tga1* alleles. Finally, we might expect field sampling of teosinte to select against plants harboring *tga1-maize*, as collectors may prioritize sampling of teosinte individuals with representative or “typical” teosinte fruitcase morphology to avoid sampling recent hybrids with maize.

Puzzlingly, we did observe *tga1-maize* in *mexicana*, which contributed to maize evolution only thousands of years after maize domestication began (Yang *et al*. 2023). Because our estimate for the age of *tga1-maize* at ≈ 41k-50kya predates the estimated divergence time between *parviglumis* and *mexicana* at ≈ 30.7kya (Chen *et al*. 2022), it is indeed possible that the allele arose in the shared common ancestor of the two subspecies. Here though, we find that gene flow from maize into *mexicana* likely accounts for our observation of *tga1-maize* in wild *mexicana*. However, this finding does not preclude the possibility that *tga1-maize* arose in *mexicana* or a shared *mexicana*/*parviglumis* ancestor. Consistent with this possibility, the two wild subspecies form multiple hybrid zones where populations exhibit varying levels of admixture (Hufnagel *et al*. 2021). Evidence indicates extensive past and ongoing gene flow between maize and *mexicana* (Ross-Ibarra et al. 2009; Yang et al. 2023). While *tga1-maize* alleles are likely selected against in *mexicana* populations (Hufford et al. 2013; Calfee et al. 2021), extensive gene flow into *mexicana* from maize continues today (Hufford et al. 2013) and our ELAI results suggest that *tga1-maize* alleles may nonetheless persist for many years.

It is often reported that the 1970s teosinte field surveys evaluated over one million teosinte plants (Iltis 2000, 2006), including populations of both *parviglumis* and *mexicana*, but failed to identify a single exposed kernel. However, the “teosinte mutant hunts” did in fact identify partially extruded kernels (Iltis 2000; Galinat 1992), but most workers presumed them to be recent backcrosses of maize-teosinte hybrids (Iltis 2000). At least some of these partially extruded kernels were grown and produced plants with wild-type fruitcases, perhaps suggesting a non-heritable cause for their maize-like phenotypes in the wild (Wilkes 2004). Alternatively, variation in genetic background and the developmental plasticity of fruitcase morphology could also explain the apparent absence of exposed kernels in the wild (Dorweiler and Doebley 1997; Wang et al. 2015). *Tga1* genotype incompletely controls fruitcase phenotypes, as the QTL containing *tga1* explains a substantial (27-62.4%) but still incomplete proportion of the phenotypic variation in measured glume traits (Doebley and Stec 1991, 1993; Briggs et al. 2007). Besides *tga1-maize* however, the genes and causal variants influencing glume induration and kernel exposure remain uncharacterized. In addition, fruitcase morphology exhibits developmental plasticity, where under some conditions the rachid and glume fail to completely enclose teosinte kernels (Piperno *et al*. 2019). Therefore, we might expect that in some genetic backgrounds and in some environments, *tga1-maize* may not strongly affect fruitcase phenotype, and the expected *tga1-maize* phenotypes may not be consistently identifiable through observational surveys alone.

### Implications for domestication

Our finding of an ancient origin for *tga1-maize* in teosinte counters decades-long assumptions concerning the fitness effects of *tga1-maize* in wild populations. Researchers long viewed the fruitcase as an adaptation that allows the kernel to survive the gastrointestinal system of seed predators. Indeed, experiments in ruminant grazers not native to teosinte’s natural range (Wilkes 1967; Cirujeda et al. 2019) find that teosinte seeds remain viable after consumption by animals. But to our knowledge, no one has experimentally evaluated the fitness effects of *tga1maize* in wild teosinte populations. The teosinte fruitcase may contribute to other functions, such as seed dormancy, an important life history trait known to evolve in response to changing environments (Willis et al. 2014) and during domestication for some crops (Fuller and Allaby 2009). Indeed, tradeoffs in dispersal, seed size, and dormancy shape seed traits across plant taxa (Venable and Brown 1988). Consistent with this possibility, germination experiments in both *parviglumis* and *mexicana* find notable differences in dormancy among populations in different environments (Avendaño López et al. 2011). We therefore argue that researchers should carefully reconsider assumptions about the selective landscape for domestication genes in crop ancestors, particularly for those where the paleoecological record remains incompletely understood. For example, it is possible that selective pressures during the late Pleistocene allowed exposed kernels to persist in *parviglumis*, but shifted along with poorly understood changes in the Mesoamerican herbivore community during this period (Ferrusquía-Villafranca *et al*. 2017). Future work would benefit from carefully planned experiments to test the fitness effects of domestication alleles and traits in crop wild relatives, ideally in multiple environments. At the same time, however, we note that modern crop wild relatives are imperfect proxies for crop ancestors, as extant wild populations have experienced their own changing landscapes and selection pressures over the intervening thousands of years.

Our age estimate for the causative mutation of ≈ 41 − 49K years ago far predates even a conservatively early estimate for the beginning of maize domestication. Maize domestication was a process likely lasting hundreds to thousands of years, but the exact timing of this period remains incompletely understood. The earliest estimate for the beginning of domestication is from (Beissinger *et al*. 2016), who estimate that maize began to diverge from teosinte populations ≈ 15, 500 years ago. However, their estimate may overestimate the onset of maize domestication due to population structure, as their teosinte sample likely represents modern populations more distantly related to maize than the one that served as the progenitor population during domestication (Beissinger *et al*. 2016). On the lower end, the earliest archaeological evidence for maize suggests that maize domestication was already underway by at least 8,700 years ago (Piperno *et al*. 2009). Not only does our estimate predate any of the current evidence for the beginning of domestication or anthropogenic selection on *tga1*, our estimate falls before the currently accepted dates for human occupation of the Americas at around 18,500-14,000 years ago (Dillehay *et al*. 2015, though see Steeves (2021) for perspectives on earlier peopling of the Americas). We acknowledge that our age estimate may not represent the precise time at which *tga1-maize* arose, given the potential for genotyping and phasing errors to affect the ability to accurately infer genealogies across the genome. However, we consistently obtain old age estimates regardless of variations in parameterization and samples included for each Relate run, increasing our confidence in our conclusions. Therefore, we argue that *tga1-maize* almost certainly arose in a wild, unmanaged teosinte population, like other key domestication loci in maize *tb1* and *gt1*, dated to *≈* 13, 000 − 28, 000 (Studer et al. 2011) and *≈*13,000 (Wills et al. 2013) years ago, respectively.

In addition to our direct age estimate of the causative allele, we argue for a standing genetic variation origin of *tga1* based on the congruence of simulated haplotype diversity patterns and mutational age distributions with those calculated from modern maize data. Our relatively simple simulations did not exhaustively explore variation in demography, selection, or recombination, and ignored genetic details such as dominance shifts, epistasis, linked selection, and genetic background. We are especially conscious that complex demographic histories may sometimes generate patterns that resemble selection on standing genetic variation (Harris et al. 2018; Garud et al. 2021). Unfortunately, the demography of ancestral teosinte populations and of maize during domestication remain poorly understood and difficult to infer, making it challenging to entirely rule out neutral processes. The simulations we do explore, however, suggest our results should be reasonably robust to many of these considerations. Nonetheless, we propose that, combined with age estimates and direct observation of *tga1-maize* in teosinte, our modeling strongly supports a pre-domestication origin of the *tga1-maize* allele.

A pre-domestication origin for *tga1-maize* prompts us to reconsider the narrative of maize domestication that has prevailed since the 1970s, when teosinte field surveys failed to produce convincing evidence for exposed kernels. For many researchers, the appearance of *tga1-maize* in cultivated or managed teosinte populations served as a “eureka” moment that initiated maize domestication (Dorweiler *et al*. 1993; Iltis 2004). Under this model, with the arrival of *tga1-maize*, early farmers finally gained access to the grain and exerted enough selective pressure —either consciously or unconsciously —that these populations began to diverge from their unmanaged relatives. Indeed, the supposed *de novo* origin of *tga1* even prompted some to consider why early cultivators would have shown interest in a plant with essentially useless grains (Smalley and Blake 2003). We argue, however, that even the earliest cultivators of teosinte would have had access to partially exposed kernels as *tga1maize* segregated in ancestral teosinte populations. Instead, as for most other crops (Larson *et al*. 2014), the catalyst for teosinte cultivation and eventually maize domestication remain up for debate.

We believe our results also have implications for archaeological interpretations of microfossils. Previous work has implicated *tga1-maize* not only in partially exposing the kernel, but also in shaping silica deposition patterns and the resulting phytolith assemblages (Dorweiler and Doebley 1997; Piperno 2009). Maize and teosinte produce distinct assemblages of phytolith, allowing archaeologists to identify maize phytoliths recovered from archaeological artifacts and sediments (Piperno 2009). In fact, the earliest evidence of archaeological maize appears not as cobs or kernels, but instead as microfossils including phytoliths (Piperno *et al*. 2009). However, whether *tga1-maize* in a teosinte genetic background produces phytolith assemblages distinct from those of maize remains unclear. If not, then teosinte with *tga1-maize* would be indistinguishable from maize in the archaeological record. Further research will be needed to understand the effects of *tga1* in different genetic backgrounds on phytolith assemblage.

Although we argue that standing genetic variation played a central role in maize domestication based on a handful of well-characterized loci, the relative importance of *de novo* vs. standing genetic variation in domestication more generally remains unanswered. Population genetic theory proposes that adaptation from standing genetic variation is expected when population sizes are large (Messer and Petrov 2013; Hermisson and Pennings 2017), the rate of beneficial mutations is high (Messer and Petrov 2013; Hermisson and Pennings 2017), and rates of outcrossing are high (Glémin and Ronfort 2013). Maize meets these expectations: ancestral teosinte populations were large (Ross-Ibarra et al. 2009; Chen et al. 2022), the relatively large genome provides many possible targets for beneficial mutations (Engelhorn *et al*. 2023), and maize and teosinte are primarily outcrossing (Hufford et al. 2011), with regional maize farming practices further promoting outcrossing in maize (Bellon et al. 2018). In contrast, *de novo* mutations may have played a more important role in rice domestication (Chen et al. 2021), consistent with its relatively small genome (Project and Sasaki 2005) and high selfing rates (Morishima and Barbier 1990). In addition to understanding how their relative contributions vary among domesticated species, further work is needed to understand how the utilization of *de novo* vs. standing genetic variation affects or interacts with domestication histories, including the timing and pace of domestication. As domestication is a useful model for understanding adaptation more broadly (Ross-Ibarra *et al*. 2007), such work will also inform our thoughts about the importance of standing variation to evolution in other taxa.

## Materials and Method

### Genomic data, filtering, and phasing

We used SNP (single nucleotide polymorphism) data from 507 maize (*Zea mays subsp. mays*), 70 *parviglumis* teosinte (*Zea mays subsp. parviglumis*), 81 *mexicana* teosinte (*Zea mays subsp. mexicana*), 14 teosinte of mixed ancestry (*Zea mays*, “mixed teosinte”), 20 *Zea diploperennis*, and 12 *Zea luxurians* individuals available as VCFs from the ZEAMAP database (Gui et al. 2020; Chen *et al*. 2022). All maize DNA sequencing data were originally acquired from the NCBI Sequence Read Archive database. The teosinte sequencing data were generated using teosinte accessions from CIMMYT, the United States Department of Agriculture, and other collaborators of (Chen *et al*. 2022), who then mapped and aligned all reads to the B73 reference version 4.0 (Jiao *et al*. 2017).

Before filtering, we first inspected the VCF at the causative site in the *tga1* locus, located at position 46350866 on chromosome 4 of the B73 reference version 4.0. We found the site to be erroneously non-biallelic, so we manually edited the VCF to remove the genotype at that site for the single teosinte individual carrying the third allele (5G3, a *Zea mays subsp. mexicana* individual) to prevent the site’s removal with downstream filtering.

We used vcftools v.0.1.13 (Danecek *et al*. 2011) to remove non-biallelic alleles (–min-alleles 2 – max-alleles 2), genotypes with genotype quality scores below 20 (–minGQ 20), and sites with missing data for more than 75% of samples (– max-missing 0.75). After filtering, we phased the VCF and imputed genotypes using beagle v.5.4 (Browning *et al*. 2021) with default settings.

### Mutational age estimates in Relate

To investigate the age of the *tga1-maize* allele, we used Relate (Speidel *et al*. 2019) to estimate the ancestral recombination graph across a roughly 5Mb region of chromosome 4 around on *tga1* (coordinates 44,000,323 to 48,999,082 on the v4 B73 maize genome). We used genotype data from (Chen *et al*. 2022) including maize inbred lines, *parviglumis, mexicana*, mixed teosinte, and 26 samples of the outgroup taxa that passed earlier quality control filtering steps, *Zea diploperennis* (n = 14) and *Zea luxurians* (n = 12). We treated the major allele across the 26 outgroup samples taxa as the ancestral state at each SNP and made an ancestral reference genome with bcftools version 1.13 (Danecek *et al*. 2021) ‘consensus’ command. We masked the reference genome with a maskfile built from shortread mapping of diverse maize to the B73 v4 reference genome (Tittes *et al*. 2021). We assumed a generation time of one year, a diploid *N*_*e*_ of 150,000, and a mutation rate of 3.3 × 10^−8^ (Clark et al. 2005).

We ran relate in parallel using the RelateParallel.sh script, and visualized the marginal tree at *tga1* with the script TreeViewMutation.sh. Our scripts and supporting files can be found on the project Github repo.

### Identification and analysis of tga1-maize in teosinte

Using bam files containing the raw reads for ZEAMAP individuals, we verified and further assessed genotypes at the causative mutation for all individuals in the ZEAMAP dataset with the SAMtools v.1.13 (Li et al. 2009) mpileup function. We used the software ELAI (Efficient Local Ancestry Inference, Guan (2014)) to estimate patterns of local ancestry in teosinte individuals heterozygous for *tga1-maize*. ELAI infers local ancestry patterns in admixed individuals by modelling patterns of linkage disequilibrium using a two-layer hidden Markov model. We selected a reference panel of 30 random tropical maize individuals, 30 *mexicana* individuals, and 30 *parviglumis* individuals to model three-way admixture. The *mexicana* and *parviglumis* individuals comprising the reference panel were previously shown to lack evidence of recent maize ancestry (Yang et al. 2023). Because ELAI models a single admixture pulse rather than ongoing gene flow, and does not simultaneously infer the time since admixture, users must designate a time since admixture. We ran ELAI with a conservatively recent time since admixture of two years. For the input, we used unphased VCFs to eliminate the potential introduction of phasing errors and retained only those SNPs that are missing in 5% or fewer individuals and with a minor allele frequency higher than 1%. We ran ELAI with 30 EM steps, 3 lower-level clusters, and 15 upper-level clusters. We separately ran three replicates of ELAI across all ten chromosomes for each individual to gain insight into genome-wide patterns of local ancestry. We verified ELAI’s ability to distinguish between maize and *mexicana* by running ELAI on two arbitrary *mexicana* individuals and one arbitrary maize individual that had been excluded from the ref12 Fairbanks and Ross-Ibarra erence panel (results not shown). We calculated the mean ELAI score (ancestry allele dosage) at each SNP from the three replicate runs per individual. We then plotted ELAI scores across chromosomes for all three heterozygous teosinte individuals. Next, we identified tracts of maize introgression in teosinte individuals as adjacent SNPs with ELAI scores larger than 0.5, representing a conservative cutoff. We combined nearby tracts fewer than 0.1 centimorgans apart, as we suspected such short gaps between tracts represent errors rather than true switches in ancestry; the length of the maize tracts at *tga1* are not qualitatively different if tracts are not combined (Table S1). To estimate the time since gene flow using the lengths of inferred tracts, we first used the linkage map for maize uplifted version 4 of the maize genome (Ogut *et al*. 2015; Calfee *et al*. 2021) to approximate the length of the maize-like region in centiMorgans (cM). We then used the equation for the expected haplotype length in cM given the number of generations (equivalent to years in maize, an annual plant) of *cM* = 100/(*g*) (Thompson 2013).

### Haplotype clustering and visualization near tga1

Before clustering, sorting, and visualizing haplotypes, we first polarized the chromosome 4 VCF to represent the SNPs as ancestral/derived, rather than reference/alternate as they were originally encoded in the ZEAMAP VCF. We polarized the chromosome 4 VCF using the polarizeVCFbyOutgroup.py script available on Kristian Ullrich’s github (Ullrich 2022). To serve as the outgroup for polarization, we generated a diploperennis consensus VCF from 14 *Zea diploperennis* samples in the ZEAMAP dataset using a custom perl script available on the project Github repo.

We used Haplostrips (Version 1.3) (Marnetto and Huerta-Sánchez 2017) to cluster and sort haplotypes at and around *tga1* (10 Kb upstream and downstream of the causative site, 4:46345866-46355866) in maize and its closest teosinte relatives (*parviglumis, mexicana*, and mixed teosinte). We implemented Haplostrips with method S 1, which clusters and sorts based on distance from a reference, for which we designated B73_1 (one of the two identical haplotypes from the inbred line B73). We used a private minor allele frequency cutoff of 0.001, which excludes SNPs present in just one haplotype. We then visualized the sorted and clustered haplotypes using a custom R script available on the project Github repo implemented in R version 4.4.2 (Team et al. 2013).

### Estimation of variable recombination rate near tga1

We ran LDhelmet (version 1.10) (Chan et al. 2012) to infer recombination rates near *tga1* in *parviglumis*. We generated the input first by subsampling the chromosome 4 VCF to include all 70 *parviglumis* individuals and subsequently to remove invariant sites with bcftools version 1.13 (Danecek et al. 2021) before subsetting a 1 Mb region centered on the causative mutation on chromosome 4 (4:45850866-46850866) with vcftools version 0.1.13 (Danecek et al. 2011). We then generated a multisample fasta using the ZmB73v4 reference fasta and the samtools (version 1.13) faidx function before using the bcftools (version 1.13) consensus function. We ran LDhelmet with default settings. Finally, we converted the output of LDhelmet from units of *ρ* per base pair to units of r (recombination events per base pair) by dividing the output by 4*N*_*e*_, where *N*_*e*_ = 100,000 for *parviglumis* (Ross-Ibarra et al. 2009). We ran LDhelmet on *parviglumis*, rather than on maize, because positive selection at *tga1* in maize likely distorts the linkage disequilibrium signals used by LDhelmet to infer local recombination rates (though (Chan et al. 2012) report that LDhelmet is more robust to effects of selection compared to other methods such as LDhat).

### Population genetic simulations of selective sweeps in msprime and SLiM Parameterization of simulations

We used msprime (version 1.2.0) and SLiM (version 4.0.1) to generate locus-scale (1 megabase) simulations of selection on *de novo* mutations and selection on standing genetic variation (Messer and Petrov 2013; Kelleher *et al*. 2018). We ran three sets of simulations with different demographic models: (1) constant-sized population with an unscaled effective population size of 100,000, (2) a simple bottleneck model adapted from Beissinger *et al*. (2016), and (3) complex bottleneck model adapted from Wang *et al*. (2017). For the Beissinger *et al*. (2016) model, we simplified the demographic model by excluding ongoing gene flow with wild teosinte. We then used the ancestral teosinte population size, bottleneck population size and timing, and exponential growth to parameterize the SLiM simulations. For the Wang *et al*. (2017) model, we converted the backwards-in-time model to a forwards-in-time model to parameterize the SLiM simulations using a custom R script available on the project Github repo. For each demographic model, we ran simulations with all combinations of five selection coefficients (unscaled 0.05, 0.01, 0.005, 0.001, and 0.0005) and two recombination rates (1e-7 and 1e-9). The recombination rates were selected from our estimates of local recombination rates at the *tga1* locus. For one approach, we used the maize linkage map derived by Ogut *et al*. (2015) and lifted to the B73 v4 reference genome by Calfee *et al*. (2021), from which we estimated per base pair recombination rates. For the second approach, we used LD-helmet to estimate local recombination rates in *parviglumis* using a linkage disequilibrium based approach (see details above). We used a mutation rate of 3 × 10^−8^ (Clark *et al*. 2005). Due to the computational limits of forward-in-time population genetic simulations and the very large population sizes for maize in the recent past, we downscaled the modeled population-sizes ten-fold. To account for the downscaled population sizes, we increased the selection coefficients, mutation rate, and recombination rates ten-fold.

Each simulation occurred in three parts. We first simulated the pre-domestication history of the population with coalescent simulations of ancestry and neutral mutations in msprime. We then simulated the domestication history of the population with forward simulations of ancestry via tree sequence tracking in SLiM. In SLiM, we modeled selective sweeps either as acting upon a *de novo* mutation, where we introduced a new mutation with a positive selection coefficient, or selection acting on a standing genetic variant, where we selected a previously neutral mutation generated in the msprime simulation and converted its neutral selection coefficient to a positive selection coefficient. For the standing variant selective sweep simulations, we tested both a “1% starting frequency” model where, at the beginning of domestication/selection, the standing variant started at a frequency of 0.08-1.12% in the population and a “5% starting frequency” model where the standing variant started at a frequency of 4.8-5.2% in the population. For both *de novo* and standing genetic variation simulations, we chose to model the selected mutation as codominant, with a dominance of h = 0.5. We have no estimate for the dominance of *tga1maize* in a teosinte background, but (Dorweiler and Doebley 1997) observe that *tga1-maize* appears partially dominant in a maize background (though do not report a dominance coefficient). Given that *tga1* appears to behave as a quantitative locus in teosinte backgrounds (Dorweiler et al. 1993), we believe that modelling *tga1-maize* as an additive allele is a reasonable assumption. We then allowed the SLiM simulation to run either until the selected mutation was lost from the population or until “present-day”, representing 15,500 years since the start of domestication. We selected 15,500 years ago as the start of domestication based on Beissinger *et al*. (2016)’s estimate of the timing of the maize population bottleneck attributed to domestication. Following the SLiM simulations, we then imported the tree sequence of the entire population into msprime, where we simplified the tree sequence, simulated neutral mutations acquired during the domestication history in SLiM, calculated mutation frequencies, output mutational ages, and generated a VCF for a subset of 500 simulated individuals per population for further analysis. Finally, we removed nonbiallelic sites from the simulated VCFs, following filtering that we performed on the empirical maize data. We ran 500 and 1000 independently seeded runs for standing genetic and *de novo* simulations, respectively.

### Analysis of simulated selective sweeps

#### Simulated mutation outcomes

We first recorded the proportion of simulation runs for each parameter set for which the mutation fixed, was lost, or was still segregating by the end of the run (representing “present-day”). For this comparison, we retained 500 random runs from each parameter set.

#### Haplotype diversity

To compare the haplotype diversity patterns of the empirical maize data and the simulated selective sweep data, we calculated haplotype homozygosity statistics H12 and H2/H1 (Garud *et al*. 2015) in the software LASSI Plus (version 1.1.1) (Harris and DeGiorgio 2020). We used a window size of 50 SNPs and a step size of 10 SNPs and the “–filter-level 2” setting, which removes monomorphic sites. H12 and H2/H1 values were calculated on the ZEAMAP chromosome 4 VCF subsetted to include just maize individuals and on each simulated VCF using the same software and settings. The H12 and H2/H1 values from 300 simulations where the mutation fixed per parameter set (demographic model, selection coefficient, recombination rate, and mutation starting frequency) were retained for further analysis. To specifically compare the patterns near the selected mutation, we used only the subset of H12 windows that included the selected mutation for both the empirical data (chr4:46350866 for *tga1*) and the simulated data. For further comparison with mutational age distributions, We then took the mean of the H2/H1 values for the sliding windows containing the selected mutation for each independently seeded simulation run.

#### Mutational age distributions

For each independent simulation run, we used tskit and pyslim (Haller and Messer 2019; Ralph *et al*. 2020; Baumdicker *et al*. 2022) to output the ages and frequency of neutral mutations within the 500 sampled simulated individuals. For a small subset of standing genetic variation simulations (*<<* 1% of all simulations), a neutral mutation arises on top of a mutation that is later selected during the domestication sweep. Due to complications with how SLiM handles such stacked mutations, we removed these simulations entirely from our analysis. For the analyzed simulations, we retained the neutral mutations located within the selective sweep determined by the start and end positions of the H12 windows. We then adjusted the mutational ages by tenfold to account for the downscaling necessary to efficiently run the simulations. For comparison to the inferred mutations ages in the observed maize data, we subset the Relate mutational age estimates to include those within the selective sweep as determined by H12/H2/H1 windows. For both simulated and observed mutational ages, we filtered to only keep mutations at intermediate frequencies (5% to 95%) in the samples.

## Data and code availability

The ZEAMAP dataset used for this project can be accessed online at https://ftp.cngb.org/pub/CNSA/data3/CNP0001565/zeamap/02_Variants/PAN_Zea_Variants/Zea-vardb/ as reported by Chen *et al*. (2022). The Github repo with analysis scripts and supporting materials will be available upon publication.

## Acknowledgments

This research was supported by funding from the National Science Foundation (award 2307175), the US Dept. of Agriculture (Hatch project CA-D-PLS-2066-H 548), and a graduate student research grant from the Center for Population Biology. We would like to thank Dolores Piperno, Leo Speidel, Elli Cryan, Michelle Stitzer, the Ross-Ibarra lab, and Graham Coop for discussions that improved the work. This research used the High-Performance Computing Core Facility (HPCCF) at the University of California, Davis. The first author would like to thank Dr. Roger Cohen of Penn Medicine, without whom this work would not have been possible. Finally, we acknowledge the smallholder farmers and indigenous peoples across the Americas, past and present, for creating and maintaining maize diversity, and whose contributions to maize research have long been overlooked.

## Supplement

**Table S1.** Length of introgressed maize tracts in teosinte individuals heterozygous for *tga1-maize*. Tracts inferred by ELAI were then converted to centiMorgans using a linkage map (Ogut *et al*. 2015) before being used to estimate the generations (years) since introgression from maize.

| sample | chromosome | start (bp) | end (bp) | start (cM) | end (cM) | length (cM) | generations |
| --- | --- | --- | --- | --- | --- | --- | --- |
| Without combining nearby tracts (0.1 cM apart) |  |  |  |  |  |  |  |
| 5A2 | 4 | 40983801 | 48275025 | 57.21 | 58.25 | 1.04 | 95.8 |
| 5G4 | 4 | 40983801 | 50470030 | 57.73 | 58.29 | 0.71 | 140.6 |
| 6A12 | 4 | 43891269 | 48667035 | 57.73 | 58.29 | 0.56 | 177.6 |
| After combining nearby tracts (0.1 cM apart) |  |  |  |  |  |  |  |
| 6A12 | 4 | 43891269 | 50010268 | 57.73 | 58.40 | 0.68 | 148.0 |

**Figure S1.**
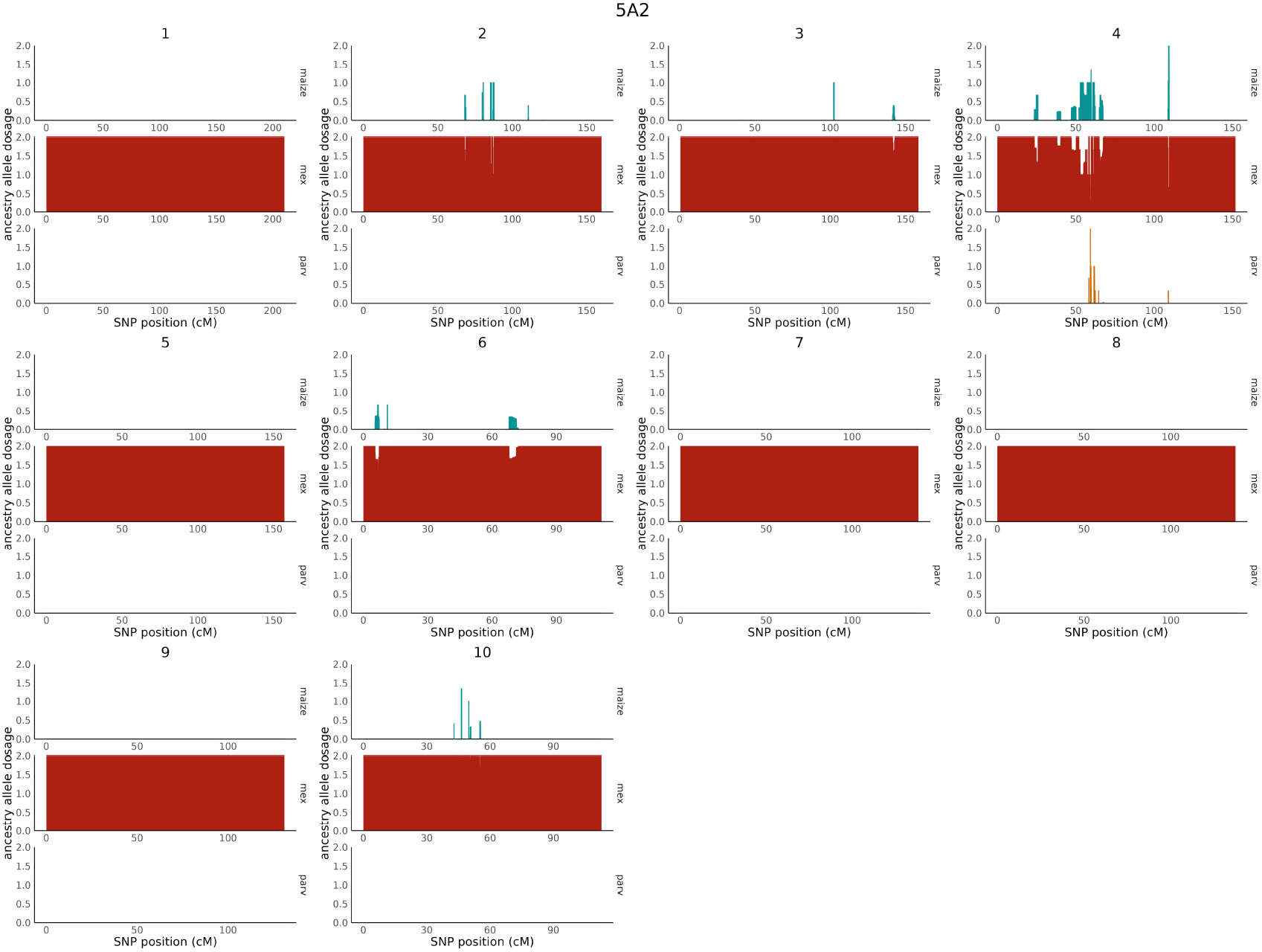
Genome-wide ancestry inferred with ELAI for *mexicana* sample 5A2. Each panel is a single chromosome, within which each ancestry origin (maize, *mexicana*, or *parviglumis*) is plotted by row.

**Figure S2.**
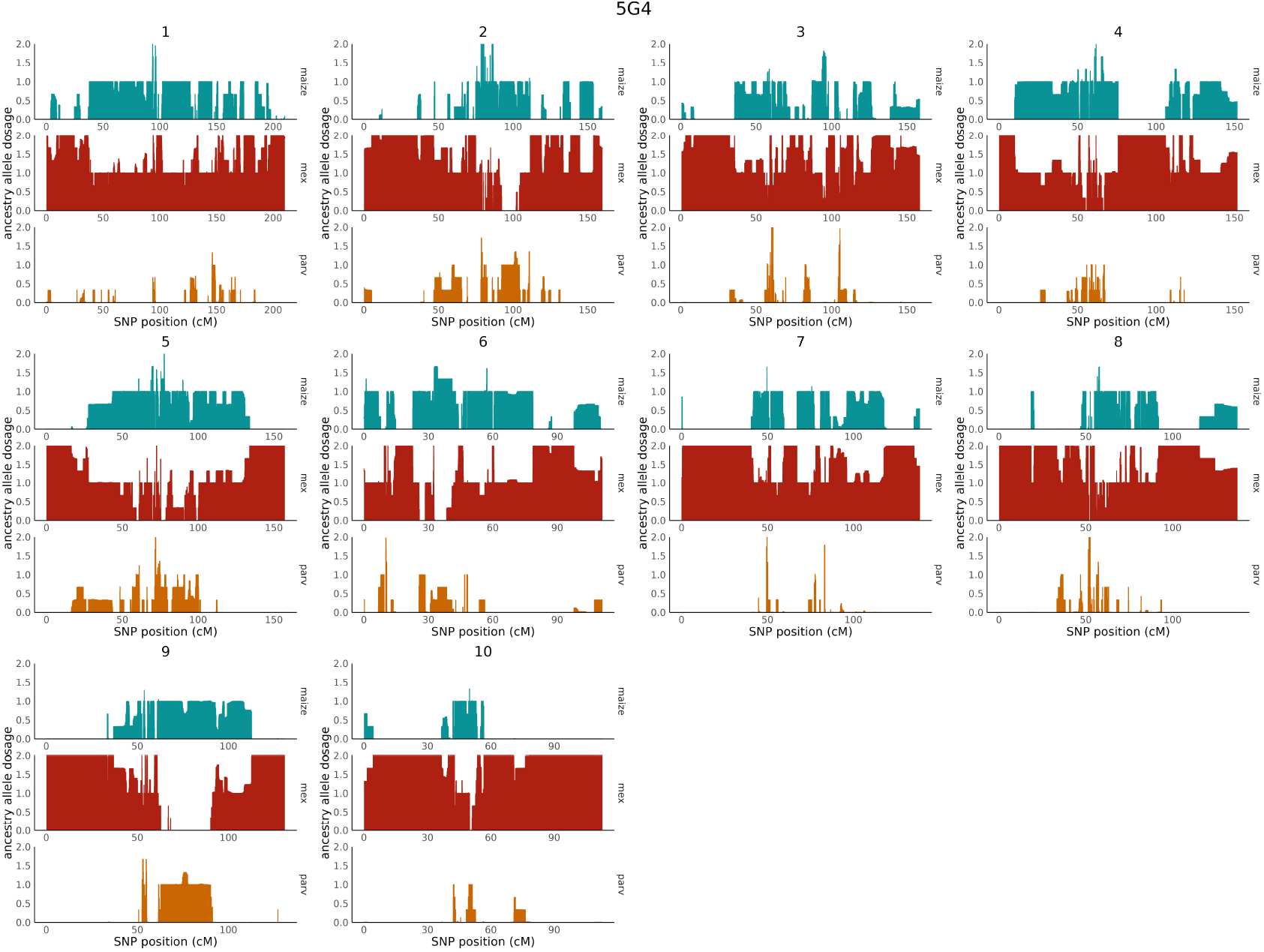
Genome-wide ancestry inferred with ELAI for mixed teosinte sample 5G4. Each panel is a single chromosome, within which each ancestry origin (maize, *mexicana*, or *parviglumis*) is plotted by row.

**Figure S3.**
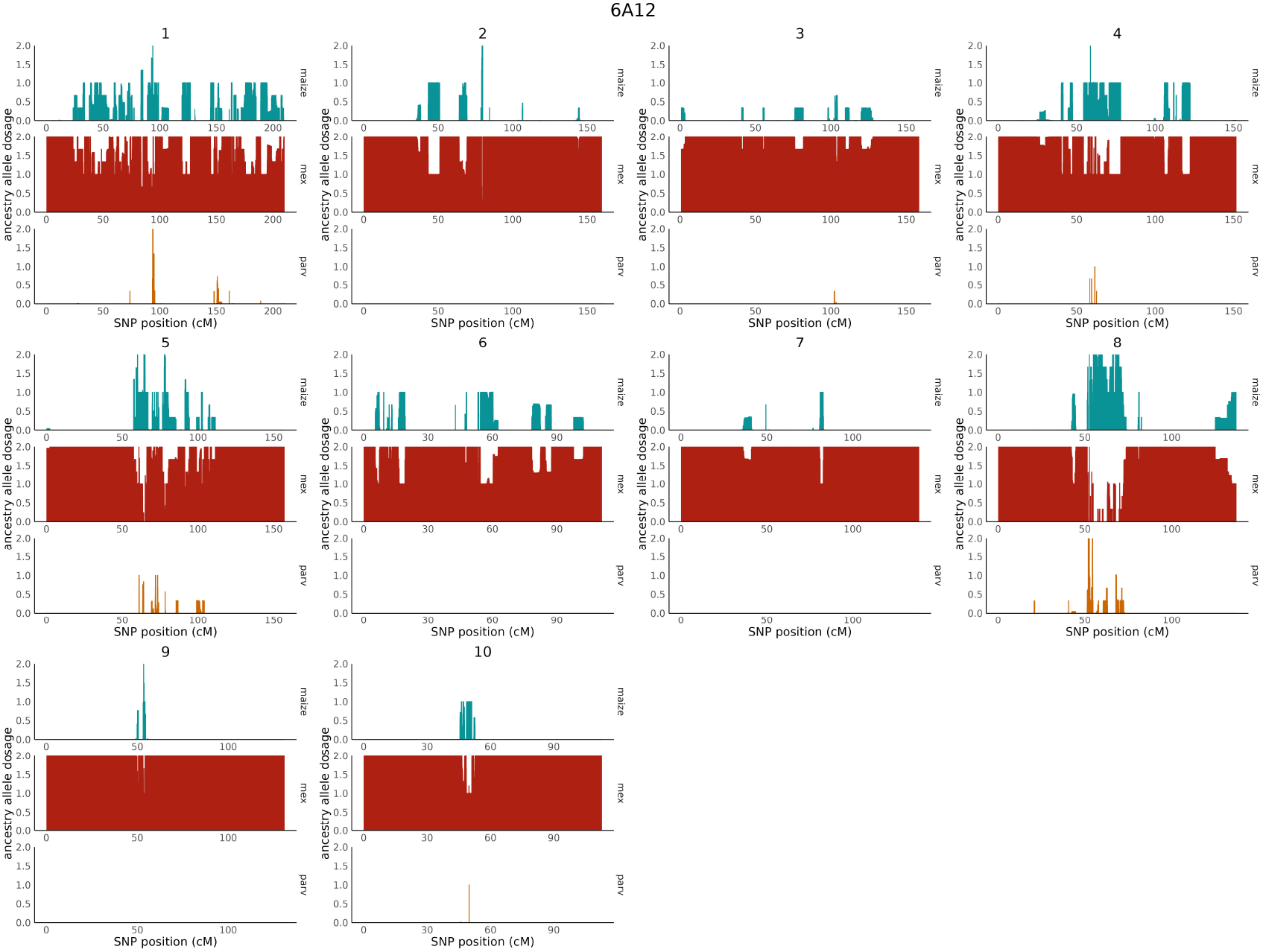
Genome-wide ancestry inferred with ELAI for *mexicana* sample 6A12. Each panel is a single chromosome, within which each ancestry origin (maize, *mexicana*, or *parviglumis*) is plotted by row.

**Figure S4.**
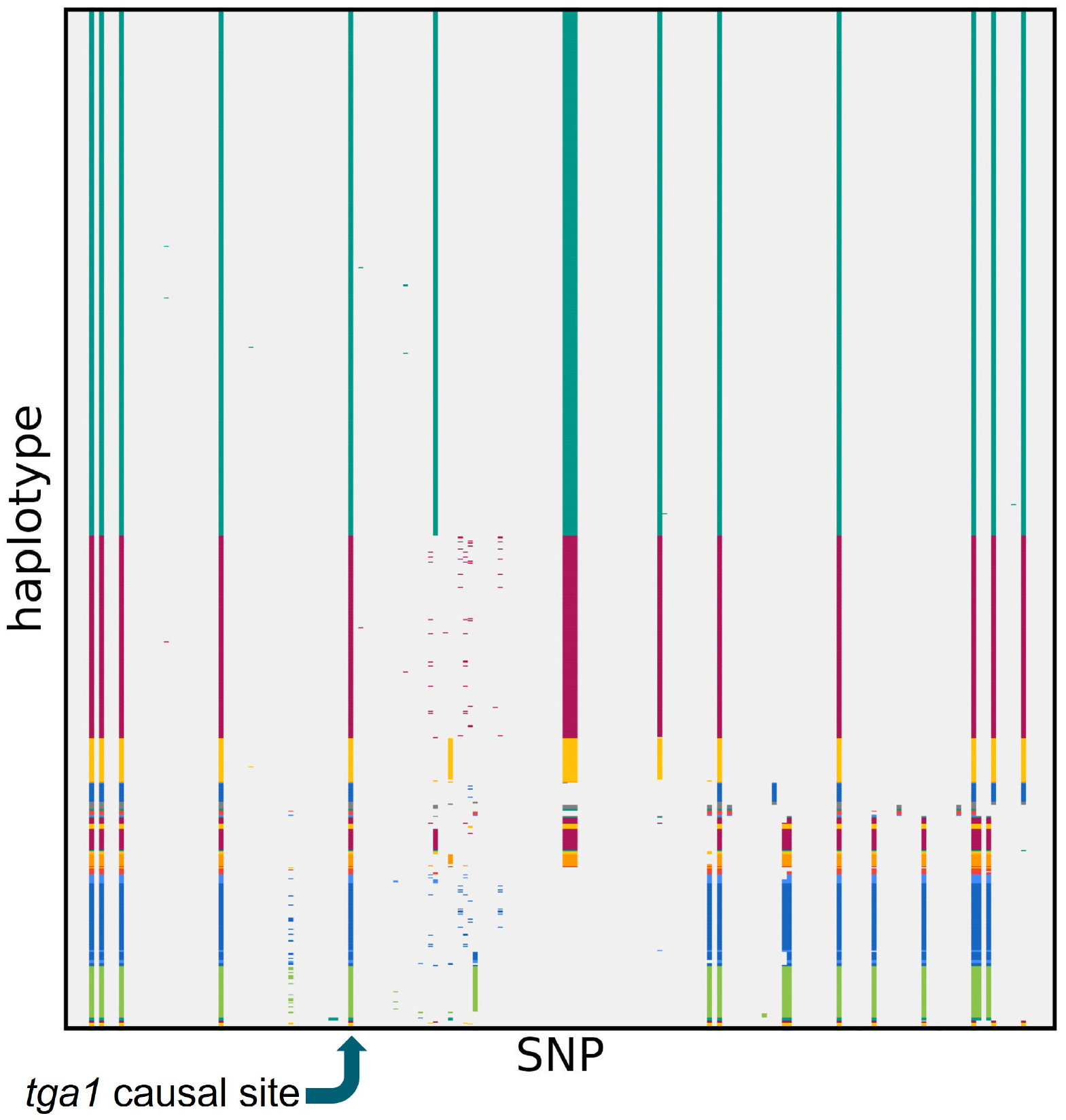
Clustered and visualized derived SNPs in a 10Kb region centered on the *tga1* causal site from 507 inbred maize lines show multiple common haplotypes present in modern maize.

**Figure S5.**
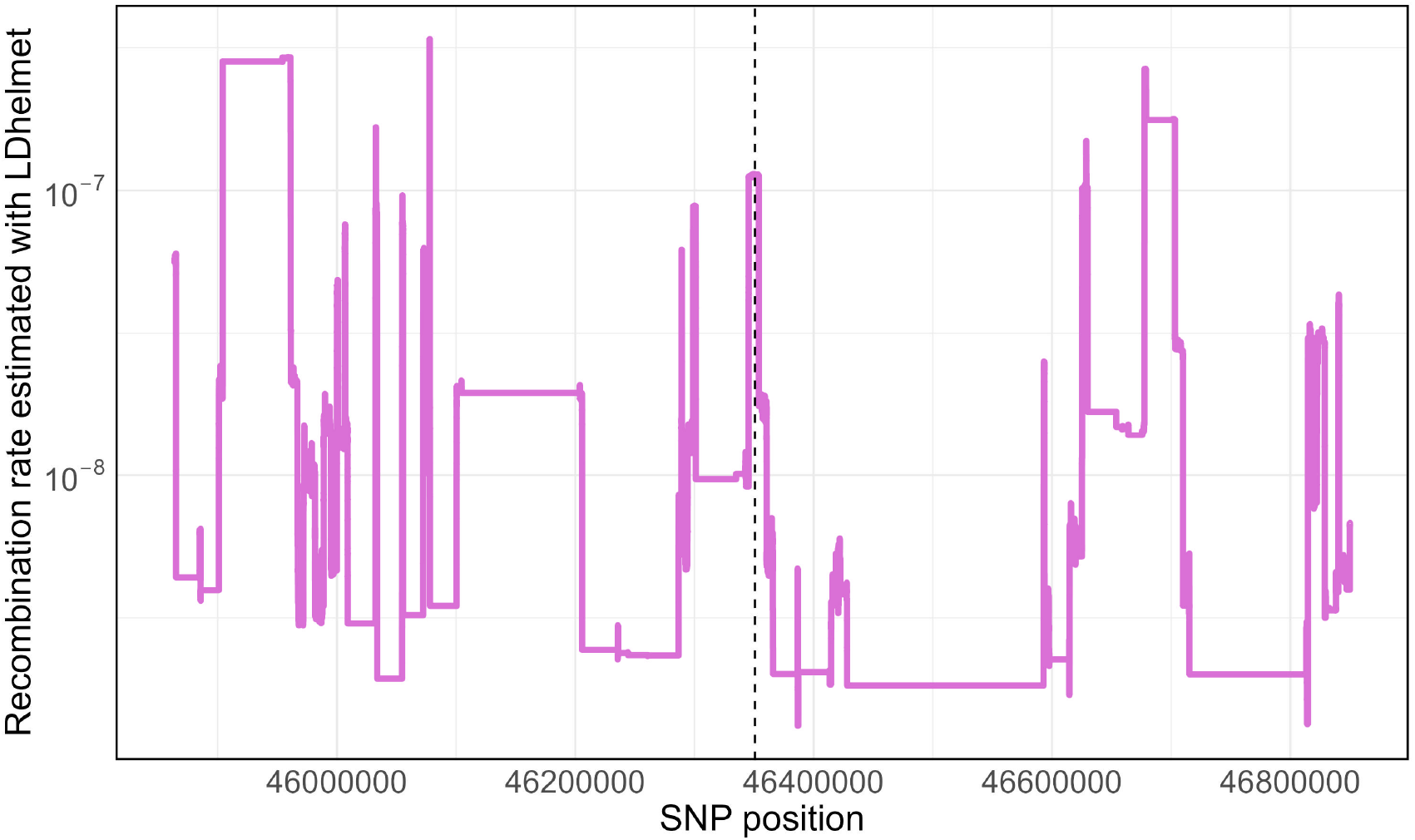
Local recombination rate inferred with LDHelmet shows elevated recombination rate at the causal *tga1* site, indicated by a dotted horizontal line. Y-axis is in log10 scale.

**Table S2.** Proportion of simulations run with the Beissinger2016 demographic model and locally elevated recombination rate (1e-7) where the mutation fixed, was lost, or was still segregating at the end of the simulation (“present-day”).

| mutation starting frequency | selection coefficient | fixed | lost | segregating |
| --- | --- | --- | --- | --- |
| 1/n | 0.05 | 0.374 | 0.626 | 0 |
| 1/n | 0.01 | 0.096 | 0.904 | 0 |
| 1/n | 0.005 | 0.056 | 0.944 | 0 |
| 1% | 0.05 | 0.998 | 0.002 | 0 |
| 1% | 0.01 | 0.71 | 0.29 | 0 |
| 1% | 0.005 | 0.528 | 0.472 | 0 |
| 5% | 0.01 | 1 | 0 | 0 |
| 5% | 0.005 | 0.962 | 0.038 | 0 |
| 5% | 0.01 | 0.966 | 0.034 | 0 |

**Table S3.** Proportion of simulations run with the Constant population size (Ne = 100,000) demographic model and locally elevated recombination rate (1e-7) where the mutation fixed, was lost, or was still segregating at the end of the simulation (“present-day”).

| mutation starting frequency | selection coefficient | fixed | lost | segregating |
| --- | --- | --- | --- | --- |
| 1/n | 0.05 | 0.364 | 0.636 | 0 |
| 1/n | 0.01 | 0.096 | 0.904 | 0 |
| 1/n | 0.005 | 0.056 | 0.944 | 0 |
| 1% | 0.05 | 1 | 0 | 0 |
| 1% | 0.01 | 1 | 0 | 0 |
| 1% | 0.005 | 1 | 0 | 0 |
| 5% | 0.05 | 1 | 0 | 0 |
| 5% | 0.01 | 1 | 0 | 0 |
| 5% | 0.005 | 1 | 0 | 0 |

**Table S4.**
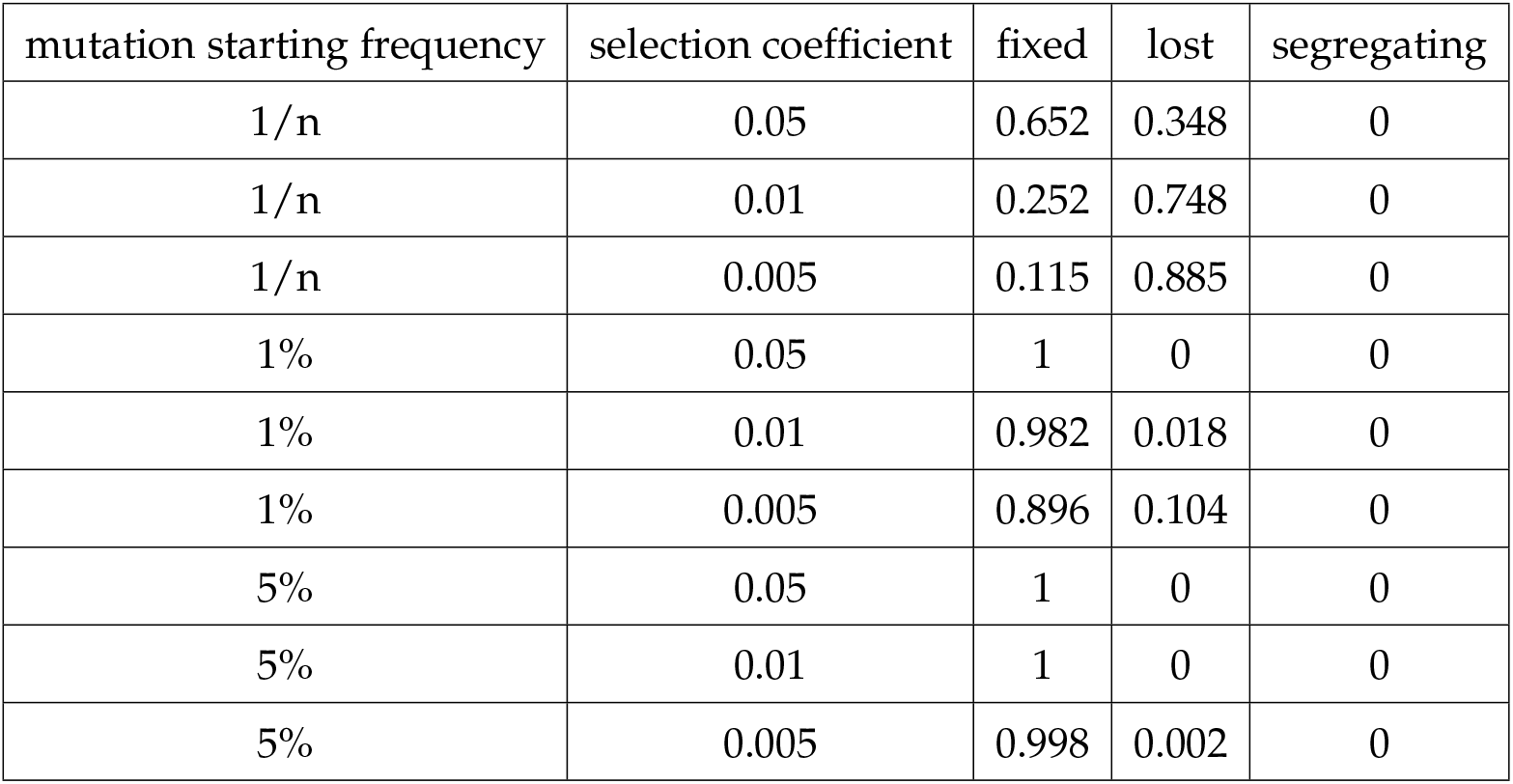
Proportion of simulations run with the Wang2017 complex bottleneck demographic model and locally elevated recombination rate (1e-7) where the mutation fixed, was lost, or was still segregating at the end of the simulation (“present-day”).

| mutation starting frequency | selection coefficient | fixed | lost | segregating |
| --- | --- | --- | --- | --- |
| 1/n | 0.05 | 0.652 | 0.348 | 0 |
| 1/n | 0.01 | 0.252 | 0.748 | 0 |
| 1/n | 0.005 | 0.115 | 0.885 | 0 |
| 1% | 0.05 | 1 | 0 | 0 |
| 1% | 0.01 | 0.982 | 0.018 | 0 |
| 1% | 0.005 | 0.896 | 0.104 | 0 |
| 5% | 0.05 | 1 | 0 | 0 |
| 5% | 0.01 | 1 | 0 | 0 |
| 5% | 0.005 | 0.998 | 0.002 | 0 |

**Figure S6.**
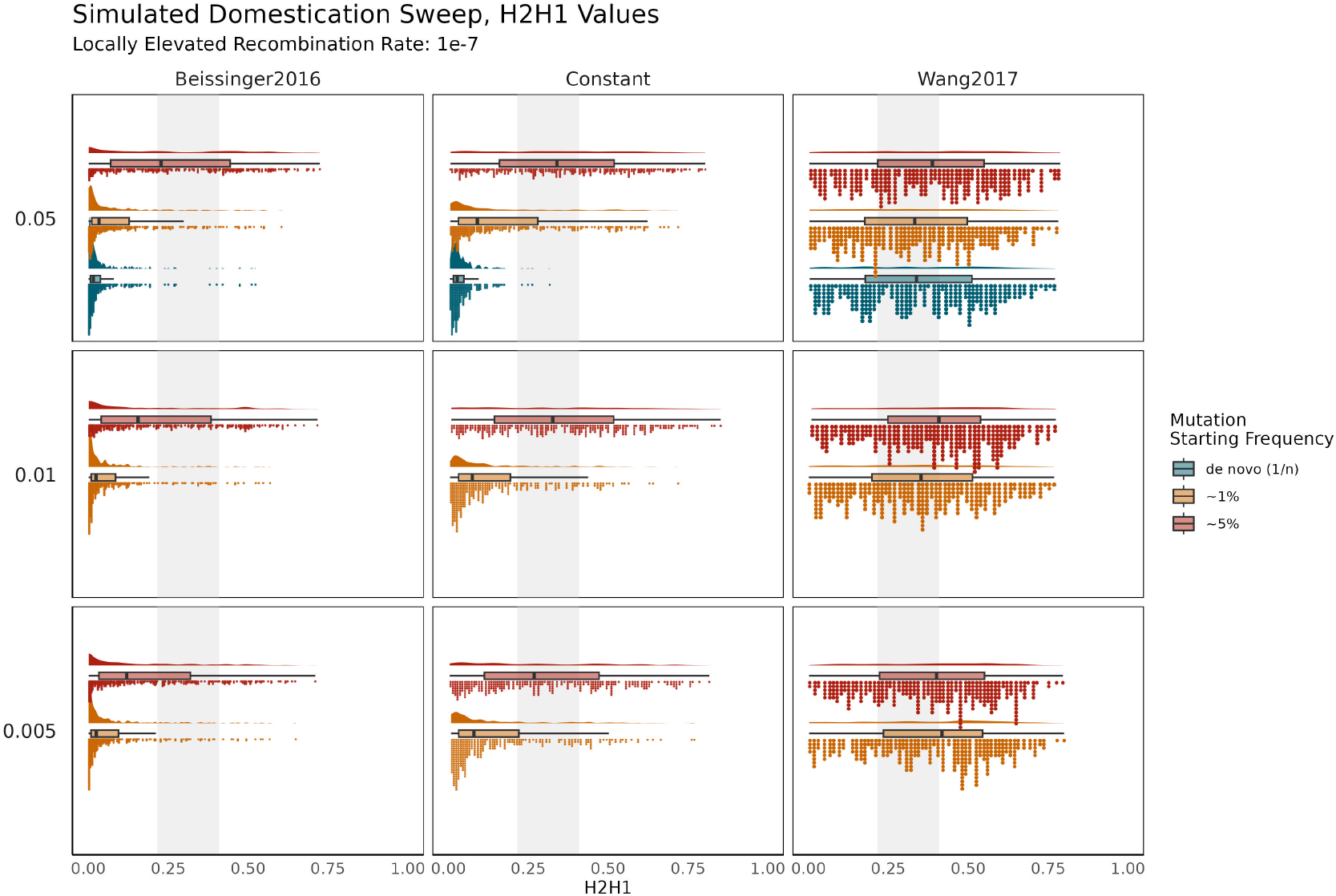
H2/H1 values from simulated sweeps under locally elevated recombination rate (1e-7) as inferred by LDHelmet. Demographic models are shown in columns, while different selection strengths are shown in rows (decreasing from top to bottom). The grey highlighted bars represent the range of observed values for 50-SNP sliding windows centered on the causal site in *tga* in a sample of 507 modern inbred maize lines.

**Figure S7.**
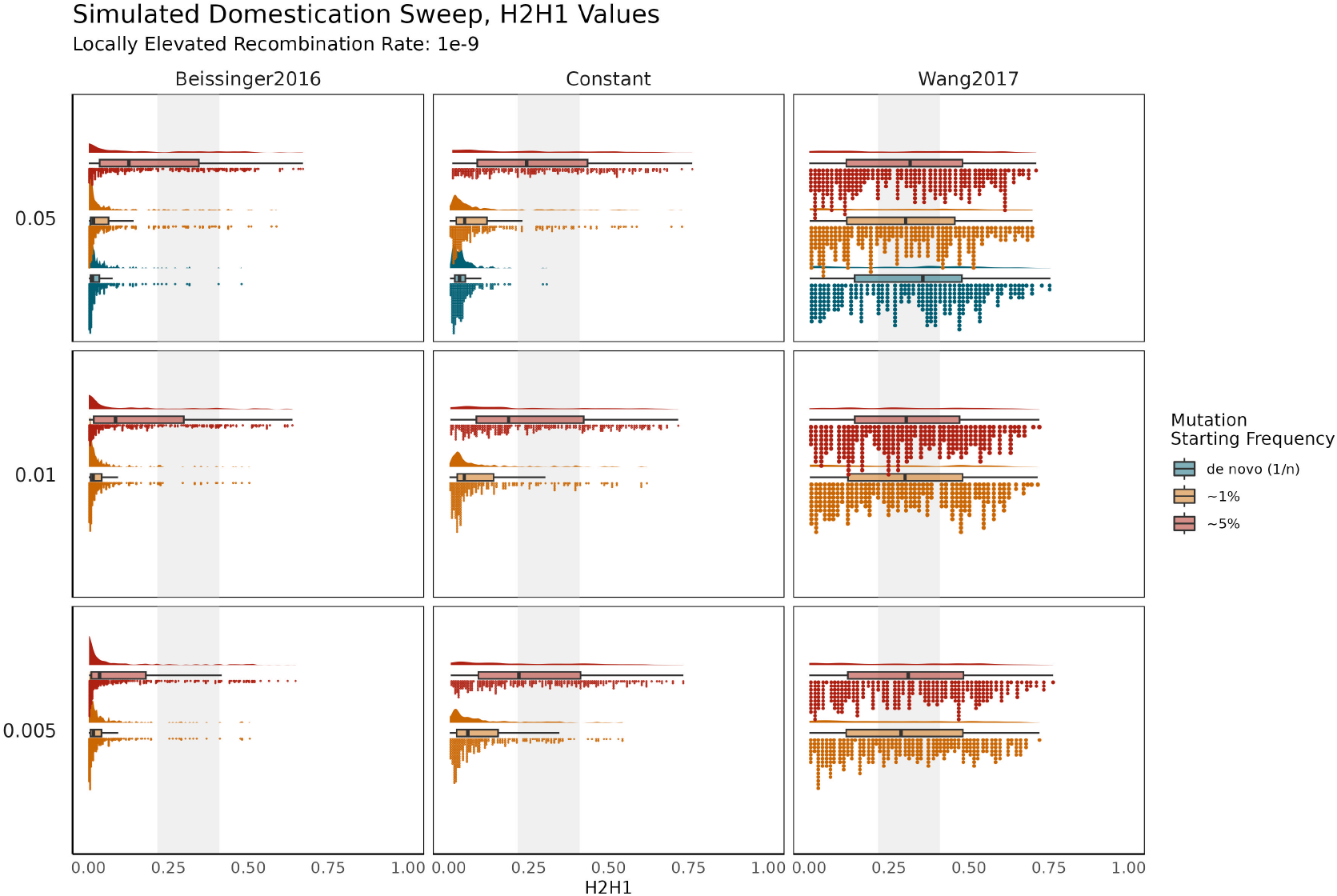
H2/H1 values from simulated sweeps under local recombination rate (1e-9) as inferred with linkage map, which is somewhat lower than similar the genome-wide average recombination rate of 8e-9. Demographic models are shown in columns, while different selection strengths are shown in rows (decreasing from top to bottom). The grey highlighted bars represent the range of observed values for 50-SNP sliding windows centered on the causal site in *tga1* in a sample of 507 modern inbred maize lines.

**Figure S8.**
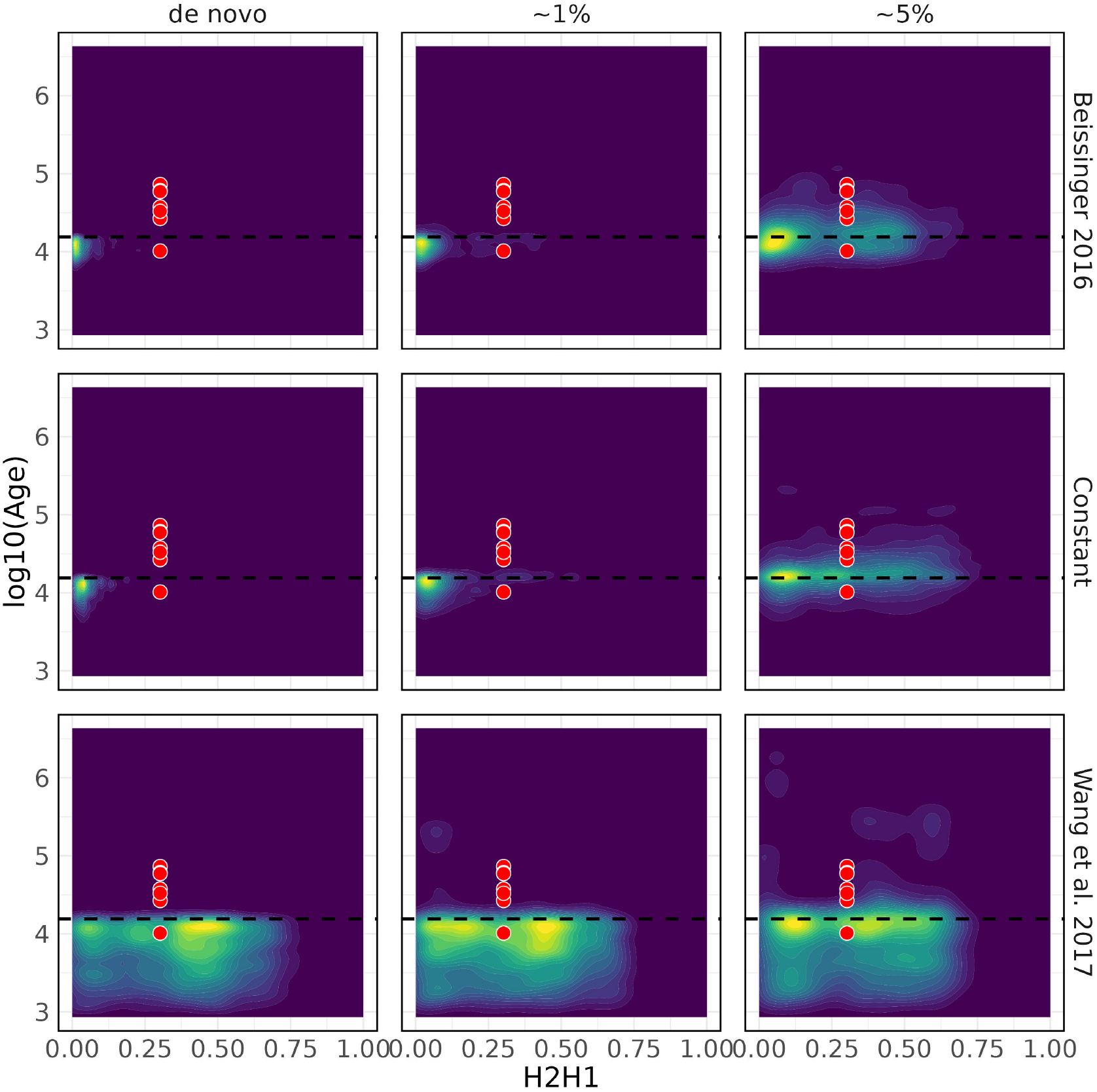
Haplotype diversity patterns (H2/H1) and mutational age distributions generated under relatively lower recombination rate generally fails to recapitulate observed patterns in modern maize at *tga1* as compared to S6, demonstrating the importance of incorporating locally elevated recombination rate into simulations. Despite the general mismatch between observed and simulated values, standing genetic variation simulations still better match observed patterns as compared to *de novo* simulations. In each plot colors range from dark blue (low density) to bright yellow (high density). Black dotted lines represent the onset of domestication (selection) at 15,500 years ago. Simulations were run with a selection coefficient of 0.05 and a recombination rate of 1e-9, estimated using the linkage map - somewhat lower than the genome-wide average of 8e-9. Red points indicate values observed in 50-SNP sliding windows centered around *tga1* in a sample of 507 modern maize inbreds.

## References

Avendaño López, A. N., J. de Jesús Sánchez González, J.A. Ruiz Corral, L. De La Cruz Larios, F. Santacruz-Ruvalcaba, et al., 2011 Seed dormancy in mexican teosinte. Crop science 51: 2056–2066.

Baumdicker, F., G. Bisschop, D. Goldstein, G. Gower, A. P. Ragsdale, et al., 2022 Efficient ancestry and mutation simulation with msprime 1.0. Genetics 220: iyab229.

Beissinger, T. M., L. Wang, K. Crosby, A. Durvasula, M. B. Hufford, et al., 2016 Recent demography drives changes in linked selection across the maize genome. Nature plants 2: 1– 7.

Bellon, M. R., A. Mastretta-Yanes, A. Ponce-Mendoza, D. Ortiz-Santamaría, O. Oliveros-Galindo, et al., 2018 Evolutionary and food supply implications of ongoing maize domestication by mexican campesinos. Proceedings of the Royal Society B 285: 20181049.

Brandenburg, J.-T., T. Mary-Huard, G. Rigaill, S. J. Hearne, H. Corti, et al., 2017 Independent introductions and admixtures have contributed to adaptation of european maize and its american counterparts. PLoS genetics 13: e1006666.

Briggs, W. H., M. D. McMullen, B. S. Gaut, and J. Doebley, 2007 Linkage mapping of domestication loci in a large maize–teosinte backcross resource. Genetics 177: 1915–1928.

Browning, B. L., X. Tian, Y. Zhou, and S. R. Browning, 2021 Fast two-stage phasing of large-scale sequence data. The American Journal of Human Genetics 108: 1880–1890.

Burke, M. K., G. Liti, and A. D. Long, 2014 Standing genetic variation drives repeatable experimental evolution in outcrossing populations of saccharomyces cerevisiae. Molecular biology and evolution 31: 3228–3239.

Calfee, E., D. Gates, A. Lorant, M. T. Perkins, G. Coop, et al., 2021 Selective sorting of ancestral introgression in maize and teosinte An ancient origin of the naked grains of maize 15 along an elevational cline. PLoS genetics 17: e1009810.

Chan, A. H., P. A. Jenkins, and Y. S. Song, 2012 Genome-wide fine-scale recombination rate variation in drosophila melanogaster. PLoS genetics 8: e1003090.

Chaturvedi, A., J. Zhou, J. A. Raeymaekers, T. Czypionka, L. Orsini, et al., 2021 Extensive standing genetic variation from a small number of founders enables rapid adaptation in daphnia. Nature Communications 12: 4306.

Chen, L., J. Luo, M. Jin, N. Yang, X. Liu, et al., 2022 Genome sequencing reveals evidence of adaptive variation in the genus zea. Nature Genetics 54: 1736–1745.

Chen, Q., W. Li, L. Tan, and F. Tian, 2021 Harnessing knowledge from maize and rice domestication for new crop breeding. Molecular Plant 14: 9–26.

Cirujeda, A., G. Pardo, A. Marí, M. Joy, and I. Casasús, 2019 Emergence and viability of teosinte seeds (zea mays ssp. mexicana ad int.) subjected to sheep digestion. Weed Research 59: 145–154.

Clark, R. M., S. Tavaré, and J. Doebley, 2005 Estimating a nucleotide substitution rate for maize from polymorphism at a major domestication locus. Molecular biology and evolution 22: 2304–2312.

Danecek, P., A. Auton, G. Abecasis, C. A. Albers, E. Banks, et al., 2011 The variant call format and vcftools. Bioinformatics 27: 2156–2158.

Danecek, P., J. K. Bonfield, J. Liddle, J. Marshall, V. Ohan, et al., 2021 Twelve years of samtools and bcftools. Gigascience 10: giab008.

Dillehay, T. D., C. Ocampo, J. Saavedra, A. O. Sawakuchi, R. M. Vega, et al., 2015 New archaeological evidence for an early human presence at monte verde, chile. PloS one 10: e0141923.

Doebley, J., 2004 The genetics of maize evolution. Annu. Rev. Genet. 38: 37–59.

Doebley, J. and A. Stec, 1991 Genetic analysis of the morphological differences between maize and teosinte. Genetics 129: 285–295.

Doebley, J. and A. Stec, 1993 Inheritance of the morphological differences between maize and teosinte: comparison of results for two f2 populations. Genetics 134: 559–570.

Dorweiler, J., A. Stec, J. Kermicle, and J. Doebley, 1993 Teosinte glume architecture 1: a genetic locus controlling a key step in maize evolution. Science 262: 233–235.

Dorweiler, J. E. and J. Doebley, 1997 Developmental analysis of teosinte glume architecture1: a key locus in the evolution of maize (poaceae). American Journal of Botany 84: 1313–1322.

Engelhorn, J., S. J. Snodgrass, A. Kok, A. S. Seetharam, M. Schneider, et al., 2023 Phenotypic variation in maize can be largely explained by genetic variation at transcription factor binding sites. bioRxiv pp. 2023–08.

Eyre-Walker, A., R. L. Gaut, H. Hilton, D. L. Feldman, and B. S. Gaut, 1998 Investigation of the bottleneck leading to the domestication of maize. Proceedings of the National Academy of Sciences 95: 4441–4446.

Ferrusquía-Villafranca, I., J. Arroyo-Cabrales, E. Johnson, J. Ruiz-González, E. Martínez-Hernández, et al., 2017 Quaternary mammals, people, and climate change: A view from southern north america. Climate change and human responses: A zooarchaeological perspective pp. 27–67.

Fukunaga, K., J. Hill, Y. Vigouroux, Y. Matsuoka, J. Sanchez G, et al., 2005 Genetic diversity and population structure of teosinte. Genetics 169: 2241–2254.

Fuller, D. Q. and R. Allaby, 2009 Seed dispersal and crop domestication: shattering, germination and seasonality in evolution under cultivation. Annual plant reviews volume 38: fruit development and seed dispersal 38: 238–295.

Galinat, W. C., 1992 Evolution of corn. Advances in Agronomy 47: 203–231.

Garud, N. R., P. W. Messer, E. O. Buzbas, and D. A. Petrov, 2015 Recent selective sweeps in north american drosophila melanogaster show signatures of soft sweeps. PLoS genetics 11: e1005004.

Garud, N. R., P. W. Messer, and D. A. Petrov, 2021 Detection of hard and soft selective sweeps from drosophila melanogaster population genomic data. PLoS Genetics 17: e1009373.

Glémin, S. and J. Ronfort, 2013 Adaptation and maladaptation in selfing and outcrossing species: new mutations versus standing variation. Evolution 67: 225–240.

Guan, Y., 2014 Detecting structure of haplotypes and local ancestry. Genetics 196: 625–642.

Gui, S., L. Yang, J. Li, J. Luo, X. Xu, et al., 2020 Zeamap, a comprehensive database adapted to the maize multi-omics era. IScience 23.

Haller, B. C. and P. W. Messer, 2019 Slim 3: forward genetic simulations beyond the wright– fisher model. Molecular biology and evolution 36: 632–637.

Haller, B. C. and P. W. Messer, 2023 Slim 4: multispecies eco-evolutionary modeling. The American Naturalist 201: E127–E139.

Harris, A. M. and M. DeGiorgio, 2020 A likelihood approach for uncovering selective sweep signatures from haplotype data. Molecular biology and evolution 37: 3023–3046.

Harris, R. B., A. Sackman, and J. D. Jensen, 2018 On the unfounded enthusiasm for soft selective sweeps ii: examining recent evidence from humans, flies, and viruses. PLoS genetics 14: e1007859.

Hermisson, J. and P. S. Pennings, 2017 Soft sweeps and beyond: understanding the patterns and probabilities of selection footprints under rapid adaptation. Methods in Ecology and Evolution 8: 700–716.

Hufford, M. B., P. Gepts, and J. ROSS-IBARRA, 2011 Influence of cryptic population structure on observed mating patterns in the wild progenitor of maize (zea mays ssp. parviglumis). Molecular ecology 20: 46–55.

Hufford, M. B., P. Lubinksy, T. Pyhäjärvi, M. T. Devengenzo, N. C. Ellstrand, et al., 2013 The genomic signature of crop-wild introgression in maize. PLoS genetics 9: e1003477.

Hufford, M. B., X. Xu, J. Van Heerwaarden, T. Pyhäjärvi, J.-M. Chia, et al., 2012 Comparative population genomics of maize domestication and improvement. Nature genetics 44: 808.

Hufnagel, D. E., K. Kananen, J. C. Glaubitz, J. de Jesuś Sánchez-González, J. F. Doebley, et al., 2021 Evidence for multiple teosinte hybrid zones in central mexico. bioRxiv.

Iltis, H., 2004 Domestication of zea: First for sugar and then for grain? a novel idea with vast implications. In 69th Annual Meeting of the Society for American Archaeology, Montreal.

Iltis, H. H., 2000 Homeotic sexual translocations and the origin of maize (zea mays, poaceae): A new look at an old problem. Economic Botany pp. 7–42.

Iltis, H. H., 2006 Origin of polystichy in maize. Multidisciplinary approaches to the prehistory, linguistics, biogeography, domestication and evolution of maiz pp. 22–54.

Jiao, Y., P. Peluso, J. Shi, T. Liang, M. C. Stitzer, et al., 2017 Improved maize reference genome with single-molecule technologies. Nature 546: 524–527.

Kelleher, J., K. R. Thornton, J. Ashander, and P. L. Ralph, 2018 Efficient pedigree recording for fast population genetics simulation. PLoS computational biology 14: e1006581.

Lai, Y.-T., C. K. Yeung, K. E. Omland, E.-L. Pang, Y. Hao, et al., 2019 Standing genetic variation as the predominant source for adaptation of a songbird. Proceedings of the National Academy of Sciences 116: 2152–2157.

Larson, G., D. R. Piperno, R. G. Allaby, M. D. Purugganan, L. Andersson, et al., 2014 Current perspectives and the future of domestication studies. Proceedings of the National Academy of Sciences 111: 6139–6146.

Li, H., B. Handsaker, A. Wysoker, T. Fennell, J. Ruan, et al., 2009 The sequence alignment/map format and samtools. bioinformatics 25: 2078–2079.

Lin, Z., X. Li, L. M. Shannon, C.-T. Yeh, M. L. Wang, et al., 2012 Parallel domestication of the shattering1 genes in cereals. Nature genetics 44: 720–724.

Liu, L., Y. Du, X. Shen, M. Li, W. Sun, et al., 2015 Krn4 controls quantitative variation in maize kernel row number. PLoS genetics 11: e1005670.

Marnetto, D. and E. Huerta-Sánchez, 2017 Haplostrips: revealing population structure through haplotype visualization. Methods in Ecology and Evolution 8: 1389–1392.

Messer, P. W. and D. A. Petrov, 2013 Population genomics of rapid adaptation by soft selective sweeps. Trends in ecology & evolution 28: 659– 669.

Moreno-Letelier, A., J. A. Aguirre-Liguori, D. Piñero, A. Vázquez-Lobo, and L. E. Eguiarte, 2020 The relevance of gene flow with wild relatives in understanding the domestication process. Royal Society Open Science 7: 191545.

Morishima, H. and P. Barbier, 1990 Mating system and genetic structure of natural populations in wild rice oryza rufipogon. Plant Species Biology 5: 31–39.

Ogut, F., Y. Bian, P. J. Bradbury, and J. B. Holland, 2015 Joint-multiple family linkage analysis predicts within-family variation better than single-family analysis of the maize nested association mapping population. Heredity 114: 552–563.

Pennings, P. S., 2012 Standing genetic variation and the evolution of drug resistance in hiv. PLoS computational biology 8: e1002527.

Peter, B. M., E. Huerta-Sanchez, and R. Nielsen, 2012 Distinguishing between selective sweeps from standing variation and from a de novo mutation. PLoS Genetics.

Piperno, D. R., 2009 Identifying crop plants with phytoliths (and starch grains) in central and south america: a review and an update of the evidence. Quaternary international 193: 146– 159.

Piperno, D. R. and K. V. Flannery, 2001 The earliest archaeological maize (zea mays l.) from highland mexico: new accelerator mass spectrometry dates and their implications. Proceedings of the National Academy of Sciences 98: 2101–2103.

Piperno, D. R., I. Holst, J. E. Moreno, and K. Winter, 2019 Experimenting with domestication: Understanding macro-and micro-phenotypes and developmental plasticity in teosinte in its ancestral pleistocene and early holocene environments. Journal of Archaeological Science 108: 104970.

Piperno, D. R., A. J. Ranere, I. Holst, J. Iriarte, and R. Dickau, 2009 Starch grain and phytolith evidence for early ninth millennium bp maize from the central balsas river valley, mexico. Proceedings of the National Academy of Sciences 106: 5019–5024.

Project, I. R. G. S. and T. Sasaki, 2005 The map-based sequence of the rice genome. Nature 436: 793–800.

Ralph, P., K. Thornton, and J. Kelleher, 2020 Efficiently summarizing relationships in large samples: a general duality between statistics of genealogies and genomes. Genetics 215: 779–797.

Ramos-Madrigal, J., B. D. Smith, J. V. Moreno-Mayar, S. Gopalakrishnan, J. Ross-Ibarra, et al., 2016 Genome sequence of a 5,310-year-old maize cob provides insights into the early stages of maize domestication. Current Biology 26: 3195–3201.

Rivera-Rodríguez, D. M., A. Mastretta-Yanes, A. Wegier, L. De la Cruz Larios, F. Santacruz-Ruvalcaba, et al., 2023 Genomic diversity and population structure of teosinte (zea spp.) and its conservation implications. Plos one 18: e0291944.

Ross-Ibarra, J., P. L. Morrell, and B. S. Gaut, 2007 Plant domestication, a unique opportunity to identify the genetic basis of adaptation. Proceedings of the National Academy of Sciences 104: 8641–8648.

Ross-Ibarra, J., M. Tenaillon, and B. S. Gaut, 2009 Historical divergence and gene flow in the genus zea. Genetics 181: 1399–1413.

Sánchez González, J. d. J., J. A. Ruiz Corral, G. M. García, G. R. Ojeda, L. D. l. C. Larios, et al., 2018 Ecogeography of teosinte. PLoS One 13: e0192676.

Sheng, Z., M. E. Pettersson, C. F. Honaker, P. B. Siegel, and Ö. Carlborg, 2015 Standing genetic variation as a major contributor to adaptation in the virginia chicken lines selection experiment. Genome biology 16: 1–12.

Smalley, J. and M. Blake, 2003 Sweet beginnings: Stalk sugar and the domestication of maize. Current Anthropology 44: 675–703.

Speidel, L., M. Forest, S. Shi, and S. R. Myers, 2019 A method for genome-wide genealogy estimation for thousands of samples. Nature genetics 51: 1321–1329.

Steeves, P. F., 2021 The indigenous Paleolithic of the western Hemisphere. U of Nebraska Press.

Studer, A., Q. Zhao, J. Ross-Ibarra, and J. Doebley, 2011 Identification of a functional transposon insertion in the maize domestication gene tb1. Nature genetics 43: 1160–1163.

Team, R. C. et al., 2013 R: A language and environment for statistical computing. Foundation for Statistical Computing, Vienna, Austria.

Tenaillon, M. I., J. U’Ren, O. Tenaillon, and B. S. Gaut, 2004 Selection versus demography: a multilocus investigation of the domestication process in maize. Molecular Biology and Evolution 21: 1214–1225.

Teotónio, H., I. M. Chelo, M. Bradić, M. R. Rose, and A. D. Long, 2009 Experimental evolution reveals natural selection on standing genetic variation. Nature genetics 41: 251–257.

Thompson, E. A., 2013 Identity by descent: variation in meiosis, across genomes, and in populations. Genetics 194: 301–326.

Tittes, S., A. Lorant, S. McGinty, J. F. Doebley, J. B. Holland, et al., 2021 Not so local: the population genetics of convergent adaptation in maize and teosinte. BioRxiv pp. 2021–09.

Ullrich, K., 2022 polarizevcfbyoutgroup.py. Accessed: 2024-11-25.

Vann, L., T. Kono, T. Pyhäjärvi, M. B. Hufford, and J. Ross-Ibarra, 2015 Natural variation in teosinte at the domestication locus teosinte branched1 (tb1). PeerJ 3: e900.

Venable, D. L. and J. S. Brown, 1988 The selective interactions of dispersal, dormancy, and seed size as adaptations for reducing risk in variable environments. The American Naturalist 131: 360–384.

Wang, H., T. Nussbaum-Wagler, B. Li, Q. Zhao, Y. Vigouroux, et al., 2005 The origin of the naked grains of maize. Nature 436: 714–719.

Wang, H., A. J. Studer, Q. Zhao, R. Meeley, and J. F. Doebley, 2015 Evidence that the origin of naked kernels during maize domestication was caused by a single amino acid substitution in tga1. Genetics 200: 965–974.

Wang, L., T. M. Beissinger, A. Lorant, C. Ross-Ibarra, J. Ross-Ibarra, et al., 2017 The interplay of demography and selection during maize domestication and expansion. Genome biology 18: 1–13.

Whipple, C. J., T. H. Kebrom, A. L. Weber, F. Yang, D. Hall, et al., 2011 grassy tillers1 promotes apical dominance in maize and responds to shade signals in the grasses. Proceedings of the National Academy of Sciences 108: E506–E512.

Wilkes, G., 2004 Corn, strange and marvelous: But is a definitive origin known. Corn: origin, history, technology, and production 4: 44.

Wilkes, H. G., 1967 Teosinte: The Closest Relative of Maize. Phd thesis, Harvard University.

Willis, C. G., C. C. Baskin, J. M. Baskin, J. R. Auld, D. L. Venable, et al., 2014 The evolution of seed dormancy: environmental cues, evolutionary hubs, and diversification of the seed plants. New Phytologist 203: 300–309.

Wills, D. M., C. J. Whipple, S. Takuno, L. E. Kursel, L. M. Shannon, et al., 2013 From many, one: genetic control of prolificacy during maize domestication. PLoS Genetics 9: e1003604.

Wright, S. I., I. V. Bi, S. G. Schroeder, M. Yamasaki, J. F. Doebley, et al., 2005 The effects of artificial selection on the maize genome. Science 308: 1310–1314.

Xu, G., X. Zhang, W. Chen, R. Zhang, Z. Li, et al., 2022 Population genomics of zea species identifies selection signatures during maize domestication and adaptation. BMC plant biology 22: 72.

Yang, N., Y. Wang, X. Liu, M. Jin, M. Vallebueno-Estrada, et al., 2023 Two teosintes made modern maize. Science 382: eadg8940.

